# *KIR2DL4* genetic diversity in a Brazilian population sample: implications for transcription regulation and protein diversity in samples with different ancestry backgrounds

**DOI:** 10.1101/2020.11.20.391649

**Authors:** Emiliana Weiss, Heloisa S. Andrade, Juliana Rodrigues Lara, Andreia S. Souza, Michelle A. Paz, Thálitta H. A. Lima, Iane O. P. Porto, Nayane dos S. B. Silva, Camila F. Bannwart Castro, Rejane M. T. Grotto, Eduardo A. Donadi, Celso T. Mendes-Junior, Erick C. Castelli

## Abstract

KIR2DL4 is an important immune modulator expressed in Natural Killer cells, being HLA-G its main ligand. We characterize *KIR2DL4* gene diversity considering the promoter, all exons, and all introns, in a highly admixed Brazilian population sample using massively parallel sequencing. We also introduce a molecular method to amplify and sequence the complete *KIR2DL4* gene. To avoid mapping bias and genotype errors commonly observed in gene families, we have developed a bioinformatic pipeline designed to minimize mapping, genotyping, and haplotyping errors. We have applied this method to survey the variability of 220 samples from the State of São Paulo, southeastern Brazil. We have also compared the *KIR2DL4* genetic diversity in Brazilian samples with the previously reported by the 1000Genomes consortium. *KIR2DL4* presents high linkage disequilibrium throughout the gene, with coding sequences associated with specific promoters. There were few, but divergent, promoter haplotypes. We have also detected many new *KIR2DL4* sequences, all with nucleotide exchanges in introns and encoding previously described proteins. Exons 3 and 4, which encode the external domains, were the most variable ones. The ancestry background influences *KIR2DL4* allele frequencies and must be considered for association studies regarding *KIR2DL4*.

## Introduction

Natural killer (NK) cells are among the major effectors in innate immunity, and they also influence adaptive immunity integrating these two complex systems. NK cells play a vital role in the immune response against infections [1] and tumors [2–4], and a series of membrane-bound receptors modulates their activity. *KIRs* (Killer Cell Immunoglobulin-Like receptors) are activating or inhibitory receptors expressed on the surface of NK cells and some T lymphocytes. They modulate NK cytotoxicity [4–6], antigen-specific cytolytic activity, and cytokine production from T lymphocytes [7].

The human *KIR* cluster is within the telomeric end of chromosome 19 and presents highly diverse haplotypes that vary in gene sequence and gene copy number [8–11]. *KIR* are transmembrane glycoproteins with two (2D) or three (3D) extracellular immunoglobulin-like (Ig-like) domains and a cytoplasmic tail, which vary in length and size. *KIR* presenting long (L) cytoplasmic domains (e.g., KIR3DL) are mostly inhibitory, and they may present one or two cytoplasmic immunoreceptor tyrosine-based inhibition motifs (ITIMs) [12]. *KIR* with a short (S) cytoplasmic domain (e.g., KIR3DS) are mostly activating receptors associated with immunoreceptor tyrosine-based activation motifs (ITAMs) and a positively charged lysine in their transmembrane region [13].

Among *KIR* genes, *KIR2DL4* is the most unusual in terms of genetic structure, expression levels, and function [5,14]. Most of the *KIR* genes are polymorphic regarding copy number. At least one copy of *KIR2DL4* locus is present in almost all *KIR* haplotypes with few exceptions [15–18], but its surface expression appears to vary among individuals. The KIR2DL4 receptor presents a single ITIM domain in its cytoplasmic tail. Depending on the ligand interacting with it, KIR2DL4 may either activate or inhibit NK cells [15].

The *KIR2DL4* region studied here comprises approximately 11 kb, considering the promoter, all exons, and all introns. There are 107 different sequences already described for this gene, encoding 54 protein variants according to the IPD-IMGT/KIR Database, *version 2.9.0* [19]. Nucleotide exchanges in the regulatory region, exons, and introns may modify gene expression patterns, and non-synonymous exchanges may influence protein structure and the affinity between ligand and the receptor. However, more than 50% of the known *KIR2DL4* alleles in the IPD-IMGT/KIR database were characterized only for exons, and there is no description of their intronic and regulatory sequences (https://www.ebi.ac.uk/ipd/kir/align.html). The *KIR2DL4* genetic diversity might be higher than we currently acknowledge.

The most studied KIR2DL4 ligands are the *HLA* class I molecules, mainly HLA-G [20,21]. HLA-G is expressed in the fetal-derived trophoblast cells modulating the immune response against fetal antigens [22–24] and in tissues such as cornea, pancreas, and thymus [25–27]. HLA-G is also expressed in pathological situations such as tumors [28–31] and autoimmune diseases [32,33]. Many nucleotide exchanges in the *KIR2DL4* locus influence its expression levels [34,35], while amino acid exchanges might disrupt the ligand-receptor binding. Those changes have been associated with recurrent abortions and preeclampsia [36–38]. However, *KIR2DL4* genetic diversity in admixed populations such as the Brazilian is unknown.

We hereby present a molecular method and a bioinformatics pipeline to evaluate *KIR2DL4*, including its distal and proximal promoter regions, all exons, and all introns, using massively parallel sequencing. We also present the *KIR2DL4* genetic diversity surveyed in 220 samples from Southeastern Brazil. Our data indicate that *KIR2DL4* is highly polymorphic among Brazilians with many new *KIR2DL4* variants. Ancestry influences allele frequencies and *KIR2DL4* presents few but divergent promoter haplotypes and a less polymorphic 3’UTR. Moreover, we demonstrate that it is mandatory to use a bioinformatics pipeline tiled for *KIR* genes to detect all *KIR2DL4* variants properly.

## Material and Methods

### Samples

We surveyed *KIR2DL4* genetic variability in 220 unrelated healthy volunteers (mean age 35, 60.97% females) from the State of São Paulo, Brazil. The Human Research Ethics Committee from the School of Medicine/Unesp has approved the study protocol (Protocol 24157413.7.0000.5411). All participants signed informed consent before blood collection. DNA was obtained by a salting out procedure and normalized to 50 ng/μL after quantification by Qubit Broad Range dsDNA Assay (Thermo Fisher Scientific Inc., Waltham, MA).

### Ancestry assessment

To estimate the ancestry contributions in this Brazilian sample, we used 34 Ancestry Informative Markers (AIM) from the SNP*for*ID 34-plex panel [39], amplified in a multiplex reaction and pooled together with the *KIR2DL4* amplicons before sequencing. We used the STRUCTURE v.2.3.4 program [40] for population structure analysis, defining K=4. For parental references, we used the 1000 Genomes data (phase 3) from Europe (404 samples from TSI, FIN, GBR, IBS populations), Africa (504 samples from YRI, LWK, GWD, MSL, and ESN populations), East Asia (504 samples from CHB, JPT, CHS, CDX, and KHV), and 14 Amerindian samples from HGDP (Surui and Karitiana) [41]. We stratified our samples within three subgroups: individuals with more than 90% of European ancestry (EUR90), individuals with more than 30% of African (AFR30), or East Asian (EAS30) ancestries. This sample represents a highly heterogeneous population sample from Brazil’s’ most populated state.

### *KIR2DL4* amplification

We designed specific primers to amplify approximately 13,387 bp of the *KIR2DL4* gene, comprising the promoter region, all exons, and all introns, in three overlapping fragments. Primers were designed in conserved regions, with no variable sites according to the *1000Genomes* project or rare variants (MAF < 0.1%) according to dbSNP version 138. These primer pairs were used to amplify *KIR2DL4* using two different approaches. For approximately half of the samples (n = 108), we amplified *KIR2DL4* separately, with no concurrent amplicons (singleplex approach). For the remaining samples (n=112), we used a multiplex approach that includes three KIR and two LILR genes, allowing us to calculate copy number differences.

### Singleplex approach

Polymerase Chain Reaction (PCR) was carried out using primers: (a) 5’-CTATTGTATGTCTTGGTGAC-3’ and 5’-CTTGATACTGGAATATTGCA-3’, which amplify the region 19:54801519-54802921 (*human genome draft assembly* hg38), producing an amplicon of 1,402 bp covering *KIR2DL4* promoter region; (b) 5’-GGGCAAACAGTGAGACCCAT-3’ and 5’-CAGGCATTTGTCCTCCCAGT-3’, which amplify the region 19:54802741-54809190 (hg38), resulting in a final product of 6,450 bp covering the first half of *KIR2DL4*; and (3) 5’-CTCTGAGATAAAACCCATTGTAA-3’ and 5’-TTCAATAAACACCTGTAAATCCC-3’ (hg38), which amplify the region 19:54807865-54814906 (hg38) resulting in an amplicon of 7,045 bp that includes the second half of *KIR2DL4*.

PCR reaction was carried out in a final volume of 25 μL. For the promoter region, we used 1.0 U of Platinum® Taq DNA polymerase and 1X buffer solution, 1.5mM of MgCl_2_, 0.20mM of each dNTP (Invitrogen-Carlsbad, CA, USA), and 0.30μM of each primer. The PCR cycling conditions were as follow: initial denaturation at 94°C for 1 minute; 32 denaturation cycles at 94°C for 30 seconds, annealing at 55°C for 30 seconds, extension at 72°C for 2 minutes; followed by a final extension at 72°C for 5 minutes and hold at 4°C. We used an alternative protocol for both amplicons of the coding region: 1.25U of TaKaRa GXL Taq® DNA polymerase and 1X buffer solution, 0.20mM of dNTP, and 0.30μM of each primer. The cycling conditions were: 30 cycles of 98°C for 10 seconds, 60°C for 15 seconds, 68°C for 7 minutes, and one hold at 4°C. For each reaction, we used 100ng of genomic DNA as a template. Amplicons were evaluated on 1% agarose gel stained with GelRed® (Biotium, Inc. Hayward, CA), quantified using Qubit dsDNA High Sensitivity Assays, normalized, pooled together, and purified using ExoSAP-IT (GE Healthcare).

### Multiplex approach

The *KIR2DL4* promoter was amplified as described in the singleplex approach. The first half of *KIR2DL4* was amplified together with the first half of *KIR3DL3*, first half of *KIR3DL2*, and the full *LILRB1* gene region. The primers for the first half of *KIR3DL3* are 5’-GGTGCATAAGGTTGGGTGTTG-3’ and 5’-GGCTTCCAGTCCTAGATCATTC – 3’, which amplify from the promoter up to intron 4. The primers for the first half of *KIR3DL2* are 5’-TTAGACACAGTTTTCTGCCCAG –3’ and 5’-GTCCCACCCCAAAAATGTCC – 3’, which amplify from the promoter up to intron 6. The primers for *LILRB1/ILT-2* are 5’-AGCGTTGAACAGGGATTGAG-3’ and 5’-GCTTGACCTAGCGATTTCAC – 3’. We have combined these primers, and the primers for the first half of *KIR2DL4*, in the following proportions: 0.6 for *KIR3DL3.* 0.56 for *KIR2DL4*, 1.04 for *LILRB1*, and 1.0 for *KIR3DL2*.

The second half of *KIR2DL4* was amplified with the second half of *KIR3DL3*, the second half of *KIR3DL2*, and the full *LILRB2/ILT4* gene region. The primers for the second half of *KIR3DL3* are 5’-GATAGACACCATGGAGGGGA-3’ and 5’-CAGATGGGGTTATGTGGACG – 3’, which amplify from intron 3 until the end of the 3’UTR. The primers for the second half of *KIR3DL2* are 5’ – GTAGCTCAAAGCATGACGTG –3’ and 5’-TACACACCACCTCACTTGTG – 3’, which amplify from intron 6 until the end of the 3’UTR. The primers for *LILRB2/ILT4* are 5’-TGGACTCATCAATCACCTACAG –3’ and 5’-AGGTTTGTGGGAGGGAGTC – 3’. We have combined these primers, and the primers for the second half of *KIR2DL4*, in the following proportions: 2.5 for *KIR3DL3*, 0.56 for *KIR2DL4*, 2.5 for *LILRB2*, 1.5 for *KIR3DL2*, and 2.0 for *KIR2DL4*.

For both reactions, PCR was carried out in a final volume of 50 μL. We used 100ng of genomic DNA as a template, 3.0U of TaKaRa GXL Taq® DNA polymerase and 1X buffer solution, 0.25mM of dNTP, and (0.80μM for the first half, 0.65 mM for the second half) of primer solution. For the first reaction (first half of each gene and *LILRB1*), the cycling conditions were: 30 cycles of 98°C for 10 seconds, 68°C for 10 minutes, and one hold at 4°C. For the second reaction (second half of each gene and *LILRB2*), the cycling conditions were: 30 cycles of 98°C for 10 seconds, 60°C for 15 seconds, 68°C for 11 minutes, and one hold at 4°C. Amplicons were evaluated on 1% agarose gel stained with GelRed® (Biotium, Inc. Hayward, CA), quantified using Qubit dsDNA High Sensitivity Assays, normalized, pooled together, and purified using ExoSAP-IT (GE Healthcare). The presence of each amplicon and consistency among reactions was evaluated by electrophoresis in Agilent BioAnalyzer DNA 12,000 kits (Agilent Technologies, CA, USA).

### *KIR2DL4* library preparation and sequencing

For library preparation, we followed recommendations for the Illumina Nextera XT Sample Preparation Kit (Illumina, Inc., San Diego, CA). Samples were multiplexed using Nextera XT Index Kit, up to 96 samples per run. However, we did not carry out the library normalization step suggested by Illumina in the Nextera manual. Instead, we quantified libraries using qPCR (Kapa Biosystems, Wilmington, MA), with a further normalization step to the recommended concentration. We used Agilent High Sensitivity DNA Kits (Agilent Technologies, CA, USA) for fragmentation pattern evaluation. Sequencing was performed using MiSeq Reagent Kit version V2 (500 cycles, 2 x 250-bp).

### Sequencing data processing

We attempted to map reads using chromosome 19 from the reference genome hg38 and the aligner BWA-MEM [42]. However, the end products of these alignments were quite unsatisfactory, with large numbers of erroneously mapped reads due to the high sequence similarity among *KIR* genes [6]. Therefore, we adopted a strategy similar to the previously used for *HLA* genes on this same sample to obtain optimized mappings [43–45]. We used hla-mapper version 3.07 (available at www.castelli-lab.net/apps/hla-mapper) but with a specific database for *KIR* genes [43]. This new database contains sequences from the IPD-IMGT/KIR [19] database and GENBANK [46] related to *KIR* genes (www.castelli-lab.net/apps/hla-mapper/).

### *KIR2DL4* copy number assessment

We have calculated the *KIR2DL4* copy number in 112 samples. These samples underwent amplification with the multiplex approach as described earlier. We calculated the mean coverage in two consecutive exons from three loci, *KIR2DL4*, *KIR3DL3*, and *LILRB1*. After, we calculated the ratios *KIR2DL4/KIR3DL3* and *KIR2DL4/LILRB1*, and used this distribution to calculate *KIR2DL4* copy number, assuming that all individuals present two copies of *KIR3DL3* and *LILRB1*. We found similar results using both *KIR3DL3* and *LILRB1* as references, except for three samples. In these cases, we noticed that *LILRB1* amplification has failed, and we used only the result from *KIR3DL3*.

### Genotype and haplotype calls

We used GATK HaplotypeCaller [47,48] (version 4.1.7) to call genotypes in the GVCF mode, concatenating all genotypes in a single VCF file using GATK GenotypeGVCFs. Variant refinement was performed using *vcfx checkpl* and *vcfx evidence* as discussed elsewhere [44]. This step ensures that only high-quality genotypes are moved forward to the haplotyping step. Variants that have not passed the vcfx filter were manually evaluated. We also annotated all variants using dbSNP version 153.

We inferred haplotypes in two steps. The first uses the GATK *ReadBackedPhasing* (RBP) routine with *phaseQualityThresh* defined as 500 to detect the phase status between neighboring heterozygous sites. RBP does not phase distant polymorphic sites, multi-allelic sites, and indels. Because of that, we used probabilistic models to get complete haplotypes considering the phase sets observed with RBP and all variants (including indels and multi-allelic sites). This second step uses a local program named phasex (*available upon request*) Phasex uses Shapeit4 [49] to infer haplotypes in bi-allelic variants taking into account the phase sets reported by RBP, in 20 independent runs. The most likely haplotypes (those in which at least 95% of the runs indicated the same haplotype) are passed forward to another round of independent runs until the number of samples with haplotypes achieving this threshold no longer increases. After that, Beagle 4.1 [50,51] infers haplotypes using bi-allelic and multi-allelic variants, also in multiple runs, and the results are compared after the final round. Finally, the most likely haplotype is passed forward as a phased VCF file. The phasex program automates all these steps.

### *KIR2DL4* allele call

The phased VCF file produced in the previous step was converted into complete sequences (exons + introns) and CDS sequences (only exons, from the first translated ATG to the stop codon) using *vcfx fasta* and vcfx transcript, respectively. We also translated each exonic sequence into proteins using Emboss transeq [52]. A local script then compared each sequence (complete, CDS, or protein) with the ones available at the IPD-IMGT/KIR Database version 2.9.0 [19], detecting whether our sequences were identical to a previously described one or new. For comparison purposes, we also evaluated *KIR2DL4* genotypes in 25 samples using PING [11].

### *KIR2DL4* on the *1000Genomes* Project dataset

To evaluate *KIR2DL4* genetic diversity in different populations, we downloaded the phased VCF data from the *1000Genomes* dataset [54], for chromosome 19 following the coordinates chr19:55312974-55326361 (GRCh37 or hg19), resulting in a total of 13,387 bp matching with the region we have sequenced. Then, we created two sequences for each individual using *vcfx fasta*.

### Other analysis

We calculated allelic and haplotype frequencies, nucleotide diversity, population differentiation, and haplotype diversity using ARLEQUIN 3.5 [53]. Linkage disequilibrium (LD) was assessed using Haploview version 4.2 [54], considering variants with minor allele frequency (MAF) over 2% and defining haplotype blocks using the confidence intervals method.

## Results

The mean ancestry composition estimates for this Brazilian population sample were 75.01% European, 16.05% African, 8.30% Asian, and 0.6% Amerindian, with individuals presenting very high proportions of European, African, or Asian ancestry and individuals with varying proportions of these three major groups.

Copy number assessment in 112 individuals analyzed using the multiplex approach indicates that 5 (4.5%) samples presented only one *KIR2DL4* copy, 105 (93.7%) presented two copies, and 2 (1.8%) presented more than two copies; complete *KIR2DL4* deletion was not observed in any sample. We also detected that both samples that carry *KIR2DL4* duplication present *KIR2DL4*0110101* and *KIR2DL4*0050101* in the same chromosome. Thus, we observed 221 copies out of the expected 224 (98.7%). When we compare allele frequencies between the samples with CNV analysis (*n* = 112) and the ones without a CNV analysis (*n* = 108), assuming two *KIR2DL4* copies for all samples in the last group, both groups present similar frequencies (exact test of population differentiation, *P* = 0.43198). Likewise, when comparing the group with CNV analysis considering the number of copies against itself but assuming two *KIR2DL4* copies for each sample, allele frequencies do not differ (*P* = 1.0000).

Because of that, and because 93.7% of the samples in the first group present two *KIR2DL4* copies, we present *KIR2DL4* SNP, allele, and allotype frequencies in three formats. The first considers only 110 individuals with one or two *KIR2DL4* copies, totalizing 215 chromosomes (five individuals have just one *KIR2DL4* copies). The second assumes that all samples present two *KIR2DL4* copies to calculate allele frequencies and SNP frequencies. The frequencies might present a small deviation from what is observed in the population. The third format illustrates the proportion of individuals presenting the reference allele for each variant, irrespectively of zygosity and copy numbers.

We detected 202 variable sites on *KIR2DL4*, as described in Table S1. Many of these variants (26.73%) occurred as singletons, and they were included in the haplotype analysis only when RBP had detected their phase status or when they were the only variable site in the sample. Some variants detected in Brazilian samples (including exonic and intronic ones), noted as A in Table S1, have not been recorded yet in the IPD-IMGT/KIR database. We have detected 22 singletons not included in the haplotyping analysis because RBP has not detected their phase status (noted as F in Table S1). Most of the variants detected in Brazil (62.9%) are not included in the 1000Genomes dataset (Table S1), even some highly frequent ones. Due to the presence of samples with only one *KIR2DL4* copy, or with two copies in the same chromosome, or even the possibility of samples presenting one chromosome with duplication and one with complete deletion, resembling a typical phenotype of one copy per chromosome, Hardy-Weinberg expectation analysis was not carried out.

In this study, we will present *KIR2DL4* genetic diversity starting from the promoter and the 5’upstream region (Table 1), followed by the *KIR2DL4* haplotypes (Tables 2 and 3), the 3’UTR haplotypes (Table 4), and lastly, the haplotypes defined for each region as extended haplotypes (Table 5), as illustrated in Figure 1.

**Fig 1.**
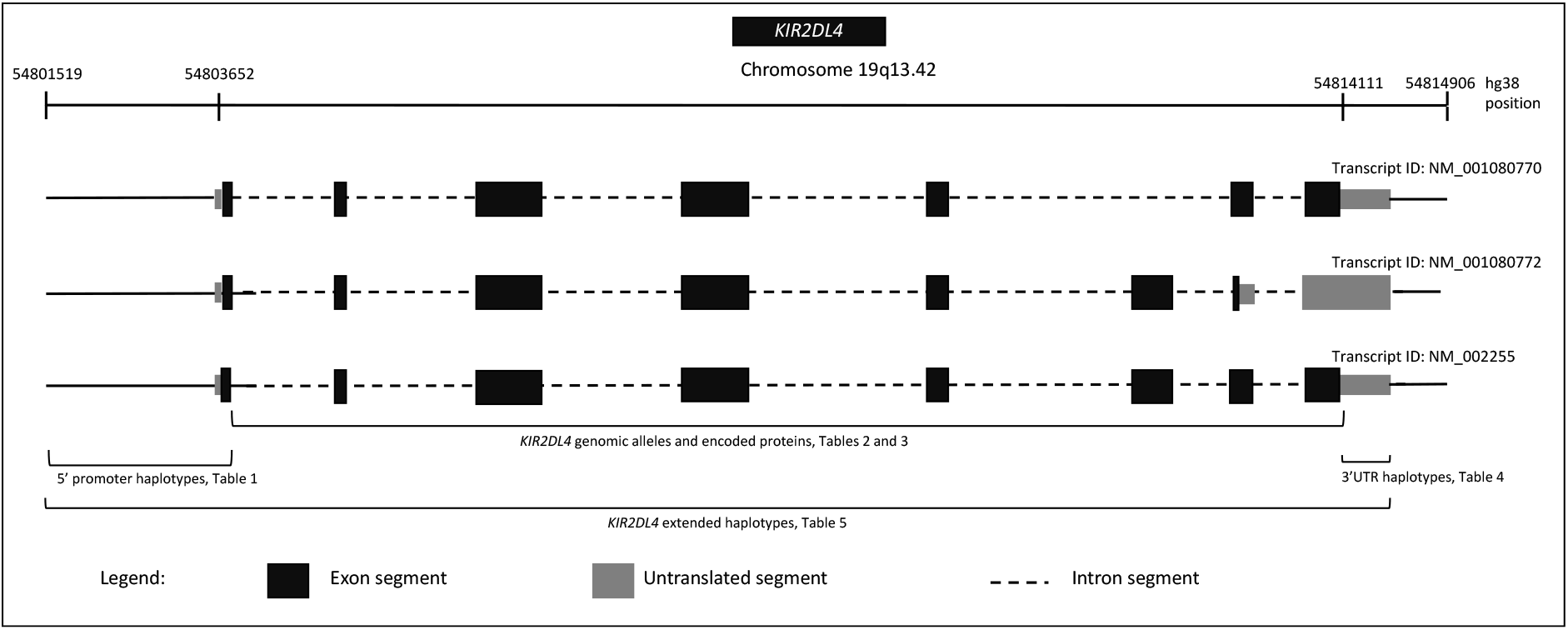
*KIR2DL4* gene structure, known transcripts, and the datasets available for each gene region.

**Table 1.** The *KIR2DL4* promoter haplotypes detected in Brazilian samples and subgroups with different ancestry backgrounds, and their frequencies.

| Haplotype Name | KIR2DL4 Promoter Haplotypes |  |  |  |  |  |  |  |  |  |  |  |  |  |  |  |  |  |  |  | Haplotype Frequencies * |  |  |  |  |  |
| --- | --- | --- | --- | --- | --- | --- | --- | --- | --- | --- | --- | --- | --- | --- | --- | --- | --- | --- | --- | --- | --- | --- | --- | --- | --- | --- |
|  | -1892 | -1545 | -1298 | -1225 | -1020 | -1013 | -973 | -820 | -507 | -362 | -321 | -306 | -302 | -273 | -270 | -229 | -227 | -215 | -210 | -158 | -46 | General <sup>a</sup><br>(2n=215) | General <sup>b</sup><br>(2n=440) | EUR90 <sup>b</sup><br>(2n = 140) | AFR30 <sup>b</sup><br>(2n = 80) | EAS30 <sup>b</sup><br>(2n = 24) |
| P01-01 | T | C | C | G | C | T | G | G | C | G | C | CA | G | T | C | C | G | G | G | T | G | 0.2279 | 0.2250 | 0.2857 | 0.2125 | 0.3333 |
| P01-02 | T | C | C | T | C | T | G | G | C | G | C | CA | G | T | C | C | G | G | G | T | G | 0.1302 | 0.1341 | 0.1500 | 0.1250 | 0.0417 |
| P01-03 | T | C | C | G | C | T | G | G | C | G | CA | CA | G | T | C | C | G | G | G | T | G | 0.0140 | 0.0068 | - | 0.0125 | - |
| P01-04 | T | C | C | T | C | T | G | G | C | G | C | CA | G | T | T | C | G | G | G | T | G | 0.0047 | 0.0045 | 0.0071 | 0.0125 | - |
| P01-05 | T | C | C | G | C | T | G | G | C | G | C | CA | G | T | C | C | G | A | G | T | G | - | 0.0023 | - | - | - |
| P02-01 | C | C | T | G | A | C | A | G | C | G | C | CA | G | T | C | C | G | G | G | T | G | 0.0791 | 0.0909 | 0.0786 | 0.0750 | 0.0833 |
| P02-02 | C | A | T | G | A | C | A | G | C | G | C | CA | G | T | C | C | G | G | G | T | G | 0.0140 | 0.0068 | - | 0.0250 | - |
| P03-01 | C | C | T | G | A | C | A | G | C | G | C | CA | G | T | C | C | G | G | C | C | G | 0.2512 | 0.2614 | 0.2429 | 0.2875 | 0.3750 |
| P03-02 | C | C | T | G | A | C | A | G | C | G | C | CA | G | T | C | T | A | G | C | C | G | 0.0186 | 0.0295 | 0.0143 | 0.0500 | - |
| P03-03 | C | C | T | G | A | C | A | G | T | G | C | CA | G | T | C | C | A | G | C | C | G | 0.0279 | 0.0182 | 0.0071 | 0.0375 | 0.0417 |
| P03-04 | C | C | T | G | A | C | A | G | C | T | C | CA | G | T | C | C | G | G | C | C | G | 0.0047 | 0.0114 | 0.0071 | 0.0125 | 0.0417 |
| P03-05 | C | C | T | G | A | C | A | G | C | G | C | CA | G | A | C | C | G | G | C | C | G | 0.0140 | 0.0114 | - | 0.0375 | - |
| P03-06 | C | C | T | G | A | C | A | G | C | G | C | CA | G | T | C | C | G | G | G | T | A | 0.0047 | 0.0023 | 0.0071 | - | - |
| P03-07 | C | C | T | G | A | C | A | G | C | G | C | CA | G | T | C | C | A | G | C | C | G | 0.0047 | 0.0023 | - | 0.0125 | - |
| P04-01 | T | C | C | G | C | T | G | A | C | G | C | C | T | T | C | C | G | G | G | T | G | 0.2047 | 0.1932 | 0.2000 | 0.1000 | 0.0833 |

a) Considering only the 110 individuals with one or two copies of *KIR2DL4*. There are 215 chromosomes because five individuals present just one *KIR2DL4* copy. b) The frequency of individuals carrying two *KIR2DL4* copies was estimated in 93.7%, one copy in 4.5%, and more than one copy in 1.8%. Because of that, to calculate these frequencies we considered all samples as presenting two *KIR2DL4* copies.
The alternative alleles regarding the human genome draft version hg38 are marked in shades of gray. Please refer to table S1 for a list of positions, SNPs, reference alleles, and alternative alleles. EUR90: Brazilian samples with at least 90% of European genomic ancestry; AFR30: Brazilians samples with at least 30% of African genomic ancestry; EAS30: Brazilians samples with at least 30% of East Asian genomic ancestry.

**Table 2.** The *KIR2DL4* alleles detected in Brazilian samples and subgroups with different ancestry backgrounds, and their frequencies.

| *KIR2DL4* alleles | Proportion of carriers<br>(*n* = 220) | General frequency<sup>a</sup><br>(2*n* = 215) | General frequency<sup>b</sup><br>(2*n* = 440) | EUR90 frequency<sup>b</sup><br>(2*n* = 140) | AFR30 frequency<sup>b</sup><br>(2*n* = 80) | EAS30 frequency<sup>b</sup><br>(2*n* = 24) |
| --- | --- | --- | --- | --- | --- | --- |
| *KIR2DL4* *0010201 | 0.2773 | 0.1535 | 0.1523 | 0.1357 | 0.1875 | 0.3333 |
| *KIR2DL4* *0010203 | 0.0273 | 0.0186 | 0.0136 | 0.0071 | 0.0125 | - |
| *KIR2DL4* *0010303 | 0.0091 | 0.0047 | 0.0045 | - | 0.0125 | - |
| *KIR2DL4* *0010304 | 0.0273 | 0.0186 | 0.0136 | 0.0071 | 0.025 | 0.0417 |
| *KIR2DL4* *0010305 | 0.0591 | 0.0279 | 0.0295 | 0.0357 | 0.0375 | - |
| *KIR2DL4* *0010306 | 0.0273 | 0.0093 | 0.0136 | 0.0071 | - | 0.0417 |
| *KIR2DL4* *0050101 | 0.2500 | 0.1488 | 0.1409 | 0.1714 | 0.125 | 0.1667 |
| *KIR2DL4* *0050102 | 0.0227 | 0.0186 | 0.0114 | 0.0214 | - | - |
| *KIR2DL4* *0050105 | 0.0045 | - | 0.0023 | 0.0071 | - | - |
| *KIR2DL4* *0050107 | 0.0091 | - | 0.0045 | 0.0143 | - | - |
| *KIR2DL4* *0060201 | 0.0455 | 0.0372 | 0.025 | 0.0214 | 0.0375 | 0.0833 |
| *KIR2DL4* *0080101 | 0.2136 | 0.1209 | 0.1182 | 0.1286 | 0.075 | 0.0833 |
| *KIR2DL4* *0080105 | 0.0136 | 0.0093 | 0.0068 | - | 0.025 | - |
| *KIR2DL4* *0080107 | 0.0773 | 0.0465 | 0.0386 | 0.05 | - | - |
| *KIR2DL4* *0080204 | 0.1727 | 0.1116 | 0.1 | 0.1143 | 0.075 | 0.0417 |
| *KIR2DL4* *0110101 | 0.1591 | 0.0698 | 0.0818 | 0.0643 | 0.1 | 0.0833 |
| Possibly new sequences | 0.3955 | 0.2046 | 0.2432 | 0.2143 | 0.2875 | 0.125 |

a) Considering only the 110 individuals with one or two copies of *KIR2DL4*. There are 215 chromosomes because five individuals present just one *KIR2DL4* copy. b) The frequency of individuals carrying two *KIR2DL4* copies was estimated in 93.7%, one copy in 4.5%, and more than one copy in 1.8%. Because of that, to calculate these frequencies we considered all samples as presenting two *KIR2DL4* copies.
Allele names were given according to the IPD-IMGT/KIR database version 2.9.0. EUR90: Brazilian samples with at least 90% of European genomic ancestry; AFR30: Brazilians samples with at least 30% of African genomic ancestry; EAS30: Brazilians samples with at least 30% of East Asian genomic ancestry.

**Table 3.** The KIR2DL4 allotypes detected in Brazilian samples and subgroups with different ancestry backgrounds, and their frequencies.

| KIR2DL4<br>allotype | Proportion of<br>carriers<br>(*n* = 220) | General<br>frequency <sup>a</sup><br>(2*n* = 215) | General<br>frequency <sup>b</sup><br>(2*n* = 440) | EUR90<br>frequency <sup>b</sup><br>(2*n* = 140) | AFR30<br>frequency <sup>b</sup><br>(2*n* = 80) | EAS30<br>frequency <sup>b</sup><br>(2*n* = 24) |
| --- | --- | --- | --- | --- | --- | --- |
| KIR2DL4*008 | 0.5273 | 0.3349 | 0.3227 | 0.3571 | 0.2000 | 0.1250 |
| KIR2DL4*001 | 0.4955 | 0.2884 | 0.3091 | 0.2571 | 0.4125 | 0.4167 |
| KIR2DL4*005 | 0.3455 | 0.2000 | 0.1977 | 0.2643 | 0.1750 | 0.1667 |
| KIR2DL4*011 | 0.1955 | 0.0930 | 0.1000 | 0.0857 | 0.1000 | 0.0833 |
| KIR2DL4*006 | 0.0500 | 0.0419 | 0.0273 | 0.0214 | 0.0375 | 0.1250 |
| KIR2DL4*013 | 0.0182 | 0.0093 | 0.0091 | - | - | 0.0417 |
| KIR2DL4*012 | 0.0136 | - | 0.0068 | - | 0.0375 | - |
| KIR2DL4*021 | 0.0091 | 0.0047 | 0.0045 | - | 0.0125 | - |
| KIR2DL4*022 | 0.0091 | 0.0047 | 0.0045 | - | - | - |
| KIR2DL4*010 | 0.0045 | 0.0047 | 0.0023 | - | 0.0125 | - |
| KIR2DL4*029 | 0.0045 | 0.0047 | 0.0023 | - | 0.0125 | - |
| Possibly new allotypes | 0.0227 | 0.0140 | 0.0136 | 0.0143 | - | 0.0417 |

a) Considering only the 110 individuals with one or two copies of *KIR2DL4*. There are 215 chromosomes because five individuals present just one *KIR2DL4* copy. b) The frequency of individuals carrying two *KIR2DL4* copies was estimated in 93.7%, one copy in 4.5%, and more than one copy in 1.8%. Because of that, to calculate these frequencies we considered all samples as presenting two *KIR2DL4* copies
Protein names (allotypes) were given according to the IPD-IMGT/KIR database version 2.9.0. EUR90: Brazilian samples with at least 90% of European genomic ancestry; AFR30: Brazilians samples with at least 30% of African genomic ancestry; EAS30: Brazilians samples with at least 30% of East Asian genomic ancestry. Some proteins only occurred in the general group (all samples together) because the individuals carrying them do not fit the subgroups.

**Table 4.** The *KIR2DL4* 3’UTR haplotypes detected in Brazilian samples and in subgroups with different ancestry backgrounds, and their frequencies.

| *KIR2DL4*<br>3'UTR haplotype | Proportion of<br>carriers<br>(*n* = 220) | General<br>frequency <sup>a</sup><br>(2*n* = 215) | General<br>frequency <sup>b</sup><br>(2*n* = 440) | EUR90<br>frequency <sup>b</sup><br>(2*n* = 140) | AFR30<br>frequency <sup>b</sup><br>(2*n* = 80) | EAS30<br>frequency <sup>b</sup><br>(2*n* = 24) | 3'UTR positions |  |  |  |  |
| --- | --- | --- | --- | --- | --- | --- | --- | --- | --- | --- | --- |
|  |  |  |  |  |  |  | 10508 | 10561 | 10597 | 10753 | 10814 |
| UTR-01 | 0.8955 | 0.7302 | 0.7386 | 0.7714 | 0.6750 | 0.5833 | G | G | T | C | C |
| UTR-02 | 0.3318 | 0.1907 | 0.1886 | 0.1643 | 0.2000 | 0.3750 | G | G | T | A | C |
| UTR-03 | 0.0909 | 0.0372 | 0.0455 | 0.0571 | 0.0500 | 0.0000 | G | C | T | C | C |
| UTR-04 | 0.0318 | 0.0233 | 0.0159 | 0.0071 | 0.0250 | 0.0417 | G | G | T | C | G |
| UTR-05 | 0.0182 | 0.0140 | 0.0091 | 0.0000 | 0.0375 | 0.0000 | A | G | T | C | C |
| UTR-06 | 0.0045 | 0.0047 | 0.0023 | 0.0000 | 0.0125 | 0.0000 | G | G | C | C | G |

a) Considering only the 110 individuals with one or two copies of *KIR2DL4*. There are 215 chromosomes because five individuals present just one *KIR2DL4* copy. b) The frequency of individuals carrying two *KIR2DL4* copies was estimated in 93.7%, one copy in 4.5%, and more than one copy in 1.8%. Because of that, to calculate these frequencies we considered all samples as presenting two *KIR2DL4* copies
The alternative alleles regarding the human genome draft version hg38 are marked in shades of gray. Please refer to table S1 for a list of positions, SNPids, reference alleles, and alternative alleles. EUR90: Brazilian samples with at least 90% of European genomic ancestry; AFR30: Brazilians samples with at least 30% of African genomic ancestry; EAS30: Brazilians samples with at least 30% of East Asian genomic ancestry.

**Table 5.** The *KIR2DL4* extended haplotypes detected in Brazilian samples, and in subgroups with different ancestry backgrounds, and their frequencies.

| Promoter Haplotype | KIR2DL4 allotype | 3'UTR Haplotype | Proportion of carriers (n = 220) | General frequency <sup>a</sup> (2n = 215) | General frequency <sup>b</sup> (2n = 440) | EUR90 frequency <sup>b</sup> (2n = 140) | AFR30 frequency <sup>b</sup> (2n = 80) | EAS30 frequency <sup>b</sup> (2n = 24) |
| --- | --- | --- | --- | --- | --- | --- | --- | --- |
| P01-01 | KIR2DL4*005 | UTR-01 | 0.3227 | 0.1814 | 0.1864 | 0.2643 | 0.1375 | 0.1667 |
| P01-01 | KIR2DL4*005 | UTR-05 | 0.0136 | 0.0140 | 0.0068 | - | 0.0375 | - |
| P01-01 | KIR2DL4*006 | UTR-01 | 0.0364 | 0.0279 | 0.0205 | 0.0214 | 0.0250 | 0.1250 |
| P01-01 | KIR2DL4*010 | UTR-01 | 0.0045 | 0.0047 | 0.0023 | - | 0.0125 | - |
| P01-01 | KIR2DL4*011 | UTR-01 | 0.0045 | - | 0.0023 | - | - | - |
| P01-01 | New protein 1 | UTR-05 | 0.0045 | - | 0.0023 | - | - | - |
| P01-01 | New protein 2 | UTR-01 | 0.0045 | - | 0.0023 | - | - | - |
| P01-01 | New protein 3 | UTR-01 | 0.0045 | - | 0.0023 | - | - | 0.0417 |
| P01-02 | KIR2DL4*008 | UTR-01 | 0.2182 | 0.1302 | 0.1273 | 0.1500 | 0.0875 | 0.0417 |
| P01-02 | KIR2DL4*012 | UTR-01 | 0.0136 | - | 0.0068 | - | 0.0375 | - |
| P01-03 | KIR2DL4*006 | UTR-01 | 0.0136 | 0.0140 | 0.0068 | - | 0.0125 | - |
| P01-04 | KIR2DL4*008 | UTR-01 | 0.0091 | 0.0047 | 0.0045 | 0.0071 | 0.0125 | - |
| P01-05 | KIR2DL4*005 | UTR-01 | 0.0045 | 0.0047 | 0.0023 | - | - | - |
| P02-01 | KIR2DL4*005 | UTR-01 | 0.0045 | - | 0.0023 | - | - | - |
| P02-01 | KIR2DL4*011 | UTR-01 | 0.1727 | 0.0744 | 0.0886 | 0.0786 | 0.0750 | 0.0833 |
| P02-02 | KIR2DL4*011 | UTR-01 | 0.0136 | 0.0140 | 0.0068 | - | 0.0250 | - |
| P03-01 | KIR2DL4*001 | UTR-01 | 0.0364 | 0.0093 | 0.0182 | 0.0071 | 0.0375 | - |
| P03-01 | KIR2DL4*001 | UTR-02 | 0.3182 | 0.1814 | 0.1795 | 0.1643 | 0.2000 | 0.3333 |
| P03-01 | KIR2DL4*001 | UTR-03 | 0.0909 | 0.0372 | 0.0455 | 0.0571 | 0.0500 | - |
| P03-01 | KIR2DL4*013 | UTR-02 | 0.0182 | 0.0093 | 0.0091 | - | - | 0.0417 |
| P03-01 | KIR2DL4*022 | UTR-01 | 0.0091 | 0.0047 | 0.0045 | - | - | - |
| P03-01 | New protein 4 | UTR-01 | 0.0045 | 0.0093 | 0.0045 | 0.0143 | - | - |
| P03-02 | KIR2DL4*001 | UTR-01 | 0.0500 | 0.0140 | 0.025 | 0.0143 | 0.0375 | - |
| P03-02 | KIR2DL4*021 | UTR-01 | 0.0091 | 0.0047 | 0.0045 | - | 0.0125 | - |
| P03-03 | KIR2DL4*001 | UTR-04 | 0.0318 | 0.0233 | 0.0159 | 0.0071 | 0.0250 | 0.0417 |
| P03-03 | KIR2DL4*001 | UTR-06 | 0.0045 | 0.0047 | 0.0023 | - | 0.0125 | - |
| P03-04 | KIR2DL4*001 | UTR-01 | 0.0227 | 0.0047 | 0.0114 | 0.0071 | 0.0125 | 0.0417 |
| P03-05 | KIR2DL4*001 | UTR-01 | 0.0227 | 0.0140 | 0.0114 | - | 0.0375 | - |
| P03-06 | KIR2DL4*011 | UTR-01 | 0.0045 | 0.0047 | 0.0023 | 0.0071 | - | - |
| P03-07 | KIR2DL4*029 | UTR-01 | 0.0045 | 0.0047 | 0.0023 | - | 0.0125 | - |
| P04-01 | KIR2DL4*008 | UTR-01 | 0.3500 | 0.2000 | 0.1909 | 0.2000 | 0.1000 | 0.0833 |
| P04-01 | New protein 4 | UTR-01 | 0.0045 | 0.0047 | 0.0023 | - | - | - |

a) Considering only the 110 individuals with one or two copies of *KIR2DL4*. There are 215 chromosomes because five individuals present just one *KIR2DL4* copy. b) The frequency of individuals carrying two *KIR2DL4* copies was estimated in 93.7%, one copy in 4.5%, and more than one copy in 1.8%. Because of that, to calculate these frequencies we considered all samples as presenting two *KIR2DL4* copies.

The *KIR2DL4* promoter presented 24 variable sites (Table S1), arranged in 15 haplotypes described in Table 1. Because the IPD-IMGT/KIR sequences do not include most of the *KIR2DL4* promoter region, the promoter haplotypes were named according to sequence similarities and the number of alternative alleles in each sequence. The most frequent haplotypes were P04-01, the reference haplotype which is identical to the sequence available in the hg38 genome draft, P01-01 and P01-02, which are one mutational step apart, and P03-01, which carries most of the alternative alleles.

The region encompassing all *KIR2DL4* exons and introns presented 178 variable sites (Table S1), arranged into 81 sequences. Some of these sequences are identical to ones already reported in the IPD-IMGT/KIR Database *version 2.9.0*, with a summed frequency of 76.8%. The frequency of each of these *KIR2DL4* alleles is in Table 2. According to the database mentioned above, the remaining sequences configure possible new *KIR2DL4* sequences, with 17 of them occurring more than once. Many of these possible new sequences are related to singletons, i.e., variants that occurred in only one sample. The frequency of some alleles is different among groups. For instance, *KIR2D4*0010201* is more frequent in the EAS30 group than EUR90 (*P* = 0.0317).

The alleles described in Table 2 encode 15 different allotypes. The most common ones were *KIR2DL4*008, KIR2DL4*001, KIR2DL4*005*, and *KIR2DL4*011.* Most of the protein sequences (eleven) have already been described, and they present a summed frequency of 98.6%. Table 3 also describes the frequency of each KIR2DL4 allotype in groups with different ancestry backgrounds. For instance, *KIR2DL4*008* is more frequent in individuals with higher European background when compared to the EAS30 group (*P* = 0.0319), *KIR2DL4*006* is frequent among individuals with higher Asian/Amerindian background (*P* = 0.0412) when compared to the EUR90 group, and *KIR2DL4*001* is less common among Europeans when compared to the AFR30 group (*P* = 0.0231).

One of the most common variants in *KIR2DL4* is an insertion/deletion at the end of exon 6 in a series of consecutive Adenines, in which 10 Adenines (10A) results in a full-length KIR2DL4 protein. In comparison, 9 Adenines (9A) leads to a premature stop codon and, consequently, to a truncated protein that lacks the ITIM motif in the cytoplasmic tail. The reference human genome (hg19 or hg38) presents the 9A allele. The truncated protein frequency in our sample (Table S1, variant rs11371265, chr6:54813220, reference allele) is 43.86%. The heterozygote proportion is 48.63%, and the homozygote proportions are 31.82% for the full-length protein and 19.54% for the truncated one. The truncated protein frequency increased among individuals with higher European (EUR90) than among higher African (AFR30) backgrounds (*P* = 0.0230).

The 3’UTR presented six different haplotypes, named UTR-01 to UTR-06, in order of frequency. Only three variable sites in the *KIR2DL4* 3’UTR region reach polymorphic frequencies, rs34785252 (UTR-02), rs376917646 (UTR-03), and rs374254587 (UTR-04 and UTR-06). The frequency of UTR-02 is higher among individuals of the EAS30 group than EUR90 (*P* = 0.0246).

We also evaluated *KIR2DL4* extended haplotypes to evaluate linkage among the promoter region, exons, and introns (Table 5). We detected many new sequences due to nucleotide exchanges in introns, but not many new exonic sequences. Table 5 lists the extended haplotypes focusing on the encoded proteins or allotypes. The promoters P01 are mostly associated with allotypes *KIR2DL4*005*, *006, *010, and *012, and the full *KIR2DL4* sequences encoding them. Promoters P02 are exclusively associated with *KIR2DL4*011.* P03 are associated with KIR2DL4*001, *021, *022, and *027, and promoter P04 is associated with *KIR2DL4*008*.

Although the number of different extended haplotypes is significant, we observed only one segregation block (Figure S1), indicating a high LD throughout *KIR2DL4*. Nucleotide diversity was 0.002580 (exons + introns), 0.002385 in the promoter region, and 0.001114 in the 3’UTR.

Comparing allele calls using our workflow and PING [11], in 25 samples, we detected a concordance of 88%. For this comparison, we used exon sequences because the current PING version evaluates only exons by default. All the divergencies were due to possibly new alleles detected by PING but not by our workflow because the genotyping of rs11371265 (the 9A/10A allele) has failed in six samples when using PING (supplementary Table S2). Nevertheless, the genotyping for all other variants when comparing both methods was identical in all samples, including a new *KIR2DL4* variant in one sample.

We compared the variants observed in the 1000Genomes Project and the ones observed in our Brazilian sample. There are 182 variable sites described in the 1000Genomes dataset in the same genome region we have evaluated, but only 75 were detected in our Brazilian samples (Table S1). Rare variants among the 1000Genomes samples (such as rs652671 and rs618871) are frequent among Brazilians. Most of the variants detected in Brazilian samples (62.9%) are not included in the 1000Genomes dataset, demonstrating that these datasets (1000Genomes and Brazilians) do not overlap. Another example is variant rs11371265 (9A/10A), which is absent from the 1000Genomes dataset, but it is described in the IPD-IMGT/KIR dataset and frequent in our Brazilian samples. We understand that these divergences between datasets are indeed a genotyping error issue.

## Discussion

Killer Immunoglobulin-like Receptors (*KIR*) are key features in modulating immune responses, and both *KIR* and their main *HLA* ligands are known to exhibit a quite substantial genetic diversity [6,8-11]. *KIR2DL4* is considered an atypical *KIR* family member in genetic structure, ligand specificity, expression levels, and signaling [55,56]. *KIR2DL4* has a central location in the *KIR* complex, and it is usually present in all individuals with rare exceptions [15-18]. Previous reports have demonstrated that NK cells and a subset of T cells (CD8+) express KIR2DL4 at different levels [55,56].

It is possible that some amino acid exchanges in both KIR2DL4 and HLA-G mature proteins might disrupt their binding, and nucleotide exchanges in introns and regulatory sequences may influence *KIR2DL4* expression patterns [57]. The complete *KIR2DL4* gene variability (including regulatory regions) has been poorly explored in real population samples, particularly admixed ones such as Brazilians. As far as we know, this is the first survey of the complete *KIR2DL4* genetic diversity in Brazilian samples, and also the first study evaluating promoter genetic diversity using NGS.

Here we investigated *KIR2DL4* genetic diversity in 220 individuals from São Paulo/Brazil. In the past, *KIR2DL4* was considered a framework gene. However, recent reports have described *KIR2DL4* deletions and duplications [11,58-61]. We have estimated that about 93.7%, 4.5%, 1.8% of our Brazilian samples present two, one, or more than two copies of *KIR2DL4*, respectively. These proportions are in agreement with previous reports [62] (allelefrequencies.net). We have calculated *KIR2DL4* allele and haplotype frequencies is three different manners. The first considers only the samples with copy number evaluation, adjusting the frequencies accordingly. The second considers that most individuals present two copies, mostly because the number of sampled copies is very close to the expected *2n* proportion. The third is the proportion of individuals carrying each allele or haplotype.

Comparing our *KIR2DL4* sequences with the ones reported on the IPD-IMGT/KIR database [19], we detected many new *KIR2DL4* alleles. The majority of these new alleles differ from the previously described ones by intronic modifications, and they encode known KIR2DL4 allotypes (Table 3). We found a few new exonic sequences. Although intronic sites are unlikely to alter gene function, modification in introns may cause alternative splicing and different expression profiles [63]. In terms of variable site number, intron 5 is the most variable one, with 67 variants, while exons 1 and 5 presented no variants (Table S1).

Because most of the variable sites lay in regulatory and intronic sequences, with only 23 variants within translated exons (Table S1), and most of the exonic nucleotide exchanges configure synonymous mutations, we have detected only 15 different KIR2DL4 allotypes. Five of them reach polymorphic frequencies, *KIR2DL4*001*, *005, *006, *008, and *011, with a summed frequency of 95.7%. These are divergent protein versions, with four amino acid exchanges in the extracellular domains. Moreover, three of these proteins encode complete KIR2DL4 molecules *(KIR2DL4*001*, *005, and *006, summed frequency of 53.41%), while *KIR2DL4*008* and *011 (summed frequency of 42.27%) encode the truncated version.

The effects of these amino acid exchanges and deletions are not clean. It is well established that *KIR2DL4* is the KIR gene with the highest sequence-identity to the ancestral KIR gene, with a direct ortholog to non-human primates such as chimpanzees [64]. The only transmembrane KIR2DL4-like sequence in orangutan ends prematurely, suggesting that this intracellular truncate motif may have a conserved biological function [65]. Although the central role of *KIR2DL4* is yet not completely understood, its evolutionary conservation suggests some impact on the survival rate. Some studies have reported that the truncated receptor is not expressed on the cell surface [34]. This could indicate that some individuals may lack functional KIR2DL4 protein expression [34,56]. However, as KIR2DL4 signals are predominantly from endosomes, unlike other *KIRs*, the fact that the truncated proteins can have a normal function should be considered. Some studies have reported that the ITIM domain does not influence the KIR2DL4 activating function through the full-length receptor [56]. Moreover, there are descriptions of fertile women lacking *KIR2DL4* [61]. These contradictions reinforce the fact that the expression and function of *KIR2DL4* remain poorly understood.

It is also unclear why some exons are highly conserved. Exon 1 was invariable both here and in the 1000Genomes dataset [66], and exon 2 presented only one rare synonymous exchange in our sample. These exons encode the leader peptide sequence. This conservation may indicate that the sequence encoded by them is essential to the correct exportation of the KIR2DL4 molecule to the cell surface and has been maintained by purifying selection. Two Ig-like domains, D0 and D2, are encoded by exons 3 and 4, respectively. Most of the variable sites within these two exons are non-synonymous exchanges (Table S1). The modification of an extracellular domain may influence the interaction between KIR2DL4 and its ligand, given that charge complementarity and protein secondary structure has a crucial role in the *KIR/HLA* recognition [67]. The linker region encoded by exon 5 was invariant in this Brazilian sample. The transmembrane regions encoded by exon 6 presented two non-synonymous exchanges, including the Adenine deletion associated with a truncated KIR2DL4 protein.

*KIR2DL4* seems to be conserved in terms of protein diversity, with only five common allotypes. This observation is compatible with previous surveys [18,34,68–71]. However, many of these studies are based on less sensitive techniques that search for known alleles only. Studies addressing *KIR* variability using modern sequencing techniques also reported low KIR2DL4 diversity at the protein level. For instance, Iranians presented only eight different exonic sequences encoding six different KIR2DL4 proteins [18], and Yupac, a small South Amerindian population, presented only three [72].

Nevertheless, few samples and different populations have been analyzed so far. The study of *KIR2DL4* genetic diversity in worldwide populations might reveal that this gene is more variable than we acknowledge, especially in intronic and regulatory regions, as we demonstrate in our Brazilian data. We found that approximately 25% of complete sequences are new when considering the promoter region, exons, and introns. However, these new sequences must be cloned and sequenced to validate the presence of new variants.

The variant rs11371265 defines either the presence of a full-length KIR2DL4 protein (the allele 10A) or a truncated version (the allele 9A), as previously reported [34,57]. Both alleles are frequent in our Brazilian sample and compatible with the observed in other populations [34,73].

However, if we take into consideration the African (AFR30) and Asian (EAS30) subgroups, the frequency of the 10A allele (the full-length protein) reaches 70% and 75%, respectively. In comparison, samples with higher European ancestry present frequencies around 54% (*P* = 0.0230 when comparing EUR90 and AFR30, but it is not significant between EUR90 and EAS30 because of the small EAS30 sample size). Thus, samples with higher African ancestry have a lower frequency of the truncated protein (about 22.5% in our sample, 28% in Congo) [68]. In another study evaluating all *KIR* genes in sub-Saharan African samples, the frequencies of *KIR2DL4*008* and *011 were very low, indicating that these populations also present a low frequency of the truncated protein [17]. Likewise, samples with higher Asian ancestry have a lower frequency of the truncated protein (about 30% in our sample, 18.9% in the Japanese population, and 19.09% in the Chinese population) [71,74]. The Amerindian population Yucpa seems to present more than 95% of full-length KIR2DL4 according to their allele frequencies [72]. The frequency of the fulllength and truncated KIR2DL4 version seems to be evenly distributed among Iranians when considering the allele frequencies [18]. Besides genetic drift, one additional explanation for these frequency shifts is selective pressure: the higher frequency of KIR2DL4 full-length proteins in Africans may somehow be related to the higher frequency of truncated *HLA-G* isoforms encoded by the *G*01:05N* allele [75,76]. Thus, more functional KIR2DL4 molecules might compensate for the lower functional HLA-G molecules in these populations.

Interestingly, the *1000Genome* phase III data do not present variant rs11371265. This phenomenon is related to common mapping bias common on paralogous genes, and it was observed for *HLA* genes [77]. Here we have developed specific tools to deal with *KIR* genes (hla-mapper + *KIR* database) and minimize mapping errors. Genomic initiatives such as the 1000Genomes project use tools to deal with the entire genome and do not focus on specific genes. Most of the variants detected in our sample (62.9%) do not overlap with 1000Genomes dataset. Therefore, publicly available data are biased for these genes, with many false-positive and falsenegative variants.

The frequency of the 9A/A10 polymorphism is influenced by ancestry. Other variants follow the same path, but our sample size only allows the detection of major differences. In this matter, *KIR2DL4*001* is highly frequent among samples with higher African ancestry, and *KIR2DL4*006* is frequent among individuals with higher Asian/Amerindian ancestry, as observed in other studies addressing samples from Asia [68,71,74]. *KIR2DL4*012* was detected only among samples with higher African background, and it is also frequent in Congo [68].

We have also characterized the complete *KIR2DL4* promoter variability. This region of approximately 2000 bases upstream the translation starting point includes the distal and proximal promoter and the 5’UTR. Previous studies indicated that *KIR2DL4* presents the most divergent promoter sequence when compared to other *KIR* genes [78]. The promoter region presents a nucleotide diversity of 0.001996, with 22 variable sites arranged into 16 haplotypes (Table 1). The core promoter (157 nucleotides upstream ATG), which is important for the transcription machinery assembly, was mostly invariable with only one rare variant at position-46.

On the other hand, *KIR2DL4* presents four divergent promoter lineages (P1, P2, P3, and P4) associated with alleles encoding different KIR2DL4 allotypes. Most of the variable sites are in absolute LD (Table 1), supporting the presence of few but divergent haplotypes. Each of these promoter lineages may define different expression profiles because of the differential binding of transcription factors. Interestingly, the major KIR2DL4 ligand, HLA-G, presents these same characteristics, i.e., four divergent promoter lineages maintained by balancing selection [79–81].

Previous studies have suggested that the first *KIR2DL4* intron could overcome the function of the promoter region when silencing elements are bond to the promoter [82]. Most of the *KIR* genes present a mini-satellite in the first intron. However, *KIR2DL4* presents a 198-bp non-repetitive sequence, which is unique among *KIR* genes [9]. Here we detected only 2 variants in this intron. Another conserved regulatory region is the 3’UTR, with only two frequent variable sites. Interestingly, the second most frequent 3’UTR haplotype, UTR-2, is more common among samples with a higher Asian background. Nevertheless, miRNA binding analyses using concordant results from miRanda and RNAhybrid indicate that both sequences present the same binding profile (data not shown).

The methodology presented here for amplification, sequencing, and data processing was suitable to detect any variable site on *KIR2DL4* promoter, exons, or introns. It relies on an algorithm implemented by the hla-mapper that optimizes alignments based on known sequences and minimizes mapping errors [43]. This method was successfully applied for classical *HLA* class I genes such as *HLA-A* and *HLA-C* [44,45]. When other genes are included in multiplex reactions, such as *KIR3DL3, LILRB1, LILRB2*, and others, we can estimate *KIR2DL4* copy number variation comparing the coverage among these loci. A similar strategy is used by PING [11]. The haplotyping method and allele calls rely on the combination of direct phasing and state-of-the-art probabilistic models (shapeit4 and Beagle). Moreover, direct phasing was obtained in 62% of the heterozygous variants, and the phasing method supports multi-allelic variants and indels. Most of the final haplotypes (summed frequency of 75%) are compatible with previously described sequences that were cloned and characterized by Sanger sequencing and deposited at the IPD-IMGT/KIR database, and this compatibility is even higher when we consider only the exonic sequences (97.0%). The new haplotypes described here are mostly due to the presence of intronic nucleotide exchanges and singletons, all presenting balanced genotypes with even proportions of reads addressing each nucleotide.

Moreover, the compatibility between the results obtained by the hla-mapper+vcfx+GATK workflow and by PING was 88% (44 out of 50 alleles, Table S2). We noticed that all the divergences were related to possible new *KIR2DL4* sequences detected by PING, mostly because genotyping of the 9A/10A variant (rs11371265) failed in six samples when using PING. No other divergencies were observed. Both methods agree with the presence of a new *KIR2DL4* variant in one sample. The hla-mapper+vcfx+GATK workflow is suitable to detect new variants, as in our Brazilian samples. As *KIR2DL4* has been poorly explored in worldwide populations, particularly in admixed populations such as Brazilians, we believe that 25% of new alleles (considering the full sequences, promoter, exons, and introns) is not surprising after an in-depth characterization such as the one performed here.

A downside of our methodology is the need of many samples to get reliable haplotypes. As we consider both read-aware phasing (using GATK) and probabilistic models to get haplotypes, the sample size is essential to get accurate haplotypes. The method is suitable to infer SNPs in a per sample model, but not haplotypes (full allele sequences). Additionally, our method is suitable to infer haplotypes in samples with one or two gene copies, but it might lose performance in samples with three or more different copies. In this case (and we have at least two samples in our dataset), we used a local Perl script to detect the best combination of alleles (available upon request). However, this Perl script does not consider regulatory sequences and introns.

Here we present *KIR2DL4* genetic diversity in regulatory regions, all exons, and all introns, in a highly admixed Brazilian population sample. We also present a molecular methodology to evaluate *KIR2DL4* using massively parallel sequencing and a bioinformatics pipeline for mapping, genotyping, haplotyping, and perform copy number calculation. *KIR2DL4* presents a high LD throughout the gene, with alleles (and their encoded proteins) associated with specific promoter sequences. There were few but divergent promoter haplotypes. The most variable exons were exons 3 and 4 (the external domains). We also describe the frequency of an insertion/deletion polymorphism in exon 6 related to a truncated KIR2DL4 protein. *KIR2DL4* allele frequencies may be influenced by sample ancestry background, indicating that ancestry must be considered for any association study regarding *KIR2DL4*.

## Data availability

The VCF file with phased genotypes is available upon request.

## Acknowledgements

This work was supported by Fundação de Amparo à Pesquisa do Estado de São Paulo – FAPESP/Brazil (Grants 2017/19223-0 and 2017/05042-4). This study was financed in part by the Coordenação de Aperfeiçoamento de Pessoal de Nível Superior (CAPES) – Finance Code 001.

---

**Fig S1.**
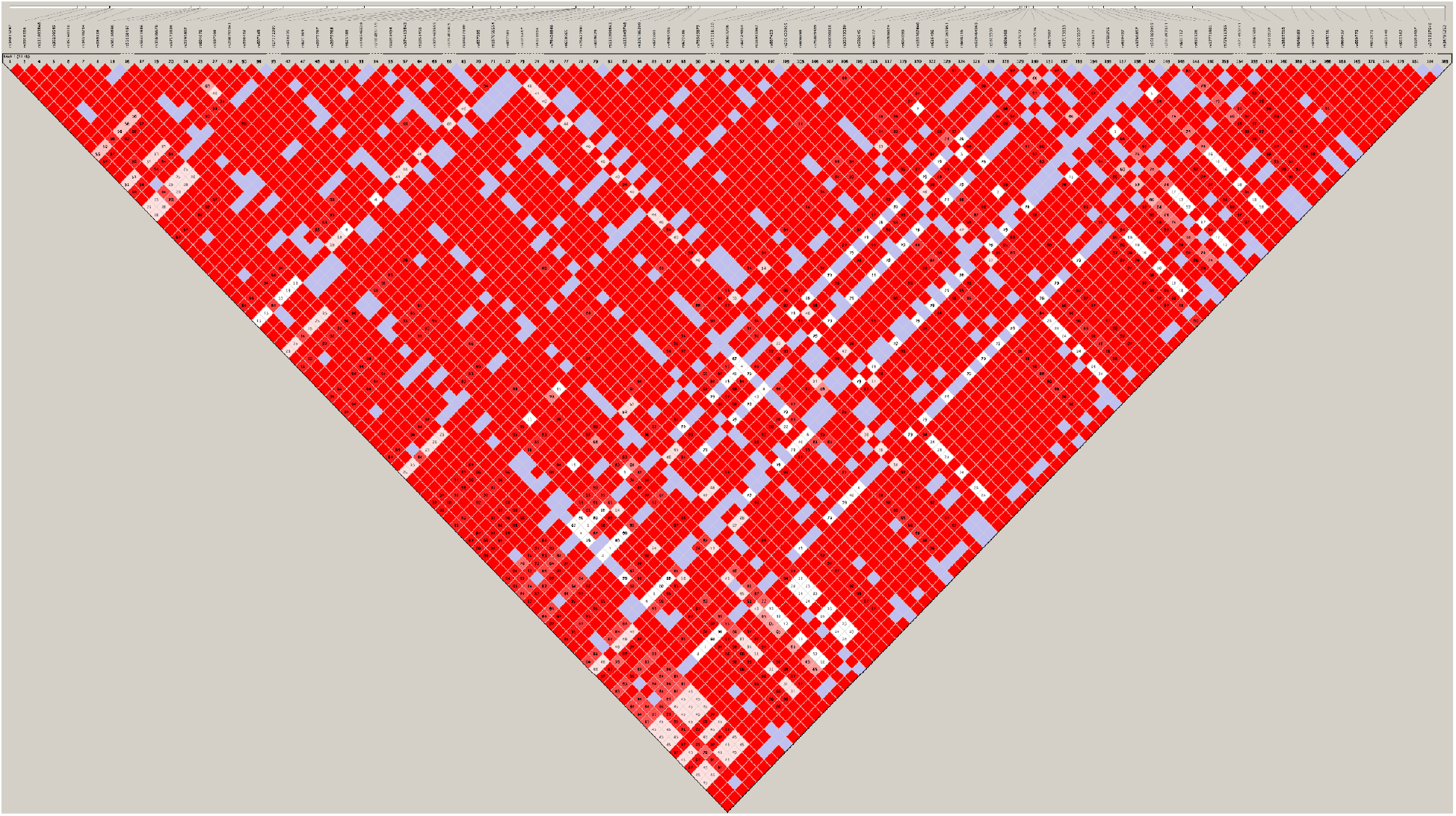
Plot of linkage disequilibrium (LD) between pairs of single nucleotide polymorphisms (SNPs), considering *KIR2DL4* variants with minimum allele frequency (MAF) > 2%, from position 19:54801519 to 19:54814906. This image was generated by Haploview 4.2 using variable sites with minimum allele frequency (MAF) > 1%. Areas in red indicate strong LD [log of odds ratio (LOD≥2, *D*’=1)]; areas in light red indicate moderate LD (LOD≥2, *D*’<1); areas in blue or almost white indicate weak LD (LOD<2, *D*’=1) and white areas indicated very weak LD (LOD<2, *D*’<1). *D*’ values different from 1 are represented inside the squares as percentages.

## REFERENCES

[1] R. Perricone, C. Perricone, C. De Carolis, Y. Shoenfeld, NK cells in autoimmunity: A two-edg’d weapon of the immune system, Autoimmun. Rev. 7 (2008) 384–390. https://doi.org/10.1016/j.autrev.2008.03.002.

[2] I. Langers, V.M. Renoux, M. Thiry, P. Delvenne, N. Jacobs, Natural killer cells: Role in local tumor growth and metastasis, Biol. Targets Ther. 6 (2012) 73–82.

[3] A.K. Purdy, K.S. Campbell, Natural killer cells and cancer: Regulation by the killer cell ig-like receptors (KIR), Cancer Biol. Ther. 8 (2009) 2209–2218. https://doi.org/10.4161/cbt.8.23.10455.

[4] S. Kumar, Natural killer cell cytotoxicity and its regulation by inhibitory receptors, Immunology. 154 (2018) 383–393. https://doi.org/10.1111/imm.12921.

[5] D. Pende, M. Falco, M. Vitale, C. Cantoni, C. Vitale, E. Munari, A. Bertaina, F. Moretta, G. Del Zotto, G. Pietra, M.C. Mingari, F. Locatelli, L. Moretta, Killer Ig-like receptors (KIRs): Their role in NK cell modulation and developments leading to their clinical exploitation, Front. Immunol. 10 (2019). https://doi.org/10.3389/fimmu.2019.01179.

[6] C. Vilches, P. Parham, KIR: Diverse, rapidly evolving receptors of innate and adaptive immunity, Annu. Rev. Immunol. 20 (2002) 217–251. https://doi.org/10.1146/annurev.immunol.20.092501.134942.

[7] L.T. van der Veken, M. Diez Campelo, M.A.W.G. van der Hoorn, R.S. Hagedoorn, H.M.E. van Egmond, J. van Bergen, R. Willemze, J.H.F. Falkenburg, M.H.M. Heemskerk, Functional Analysis of Killer Ig-Like Receptor-Expressing Cytomegalovirus-Specific CD8 + T Cells, J. Immunol. 182 (2009) 92–101. https://doi.org/10.4049/jimmunol.182.1.92.

[8] M. Uhrberg, N.M. Valiante, B.P. Shum, H.G. Shilling, K. Lienert-Weidenbach, B. Corliss, D. Tyan, L.L. Lanier, P. Parham, Human Diversity in Killer Cell Inhibitory Receptor Genes et al Human NK cells use two types of structure as their, Immunity. 7 (1997) 753–763. https://ac.els-cdn.com/S1074761300803945/1-s2.0-S1074761300803945-main.pdf?_tid=589c76df-03cf-497c-b31b-e7737f4ae42e&acdnat=1539789168_f874d148913f78499a980240ceb1d183.

[9] J. Trowsdale, R. Barten, A. Haude, C. Andrew Stewart, S. Beck, M.J. Wilson, The genomic context of natural killer receptor extended gene families, Immunol. Rev. 181 (2001) 20–38. https://doi.org/10.1034/j.1600-065X.2001.1810102.x.

[10] I. Wagner, D. Schefzyk, J. Pruschke, G. Schöfl, B. Schöne, N. Gruber, K. Lang, J. Hofmann, C. Gnahm, B. Heyn, W.M. Marin, R. Dandekar, J.A. Hollenbach, J. Schetelig, J. Pingel, P.J. Norman, J. Sauter, A.H. Schmidt, V. Lange, Allele-Level KIR Genotyping of More Than a Million Samples: Workflow, Algorithm, and Observations, Front. Immunol. 9 (2018) 2843. https://doi.org/10.3389/fimmu.2018.02843.

[11] P.J. Norman, J.A. Hollenbach, N. Nemat-Gorgani, W.M. Marin, S.J. Norberg, E. Ashouri, J. Jayaraman, E.E. Wroblewski, J. Trowsdale, R. Rajalingam, J.R. Oksenberg, J. Chiaroni, L.A. Guethlein, J.A. Traherne, M. Ronaghi, P. Parham, Defining KIR and HLA Class I Genotypes at Highest Resolution via High-Throughput Sequencing, Am. J. Hum. Genet. 99 (2016) 375–391. https://doi.org/10.1016/j.ajhg.2016.06.023.

[12] E. Vivier, M. Daëron, Immunoreceptor tyrosine-based inhibition motifs, Immunol. Today. (1997). https://doi.org/10.1016/S0167-5699(97)80025-4.

[13] L.L. Lanier, B.C. Cortiss, J. Wu, C. Leong, J.H. Phillips, Immunoreceptor DAP12 bearing a tyrosine-based activation motif is involved in activating NK cells, Nature. (1998). https://doi.org/10.1038/35642.

[14] P. Parham, MHC class I molecules and KIRS in human history, health and survival, Nat. Rev. Immunol. 5 (2005) 201–214. https://doi.org/10.1038/nri1570.

[15] A.M. Martin, E.M. Freitas, C.S. Witt, F.T. Christiansen, The genomic organization and evolution of the natural killer immunoglobulin-like receptor (KIR) gene cluster, Immunogenetics. 51 (2000) 268–280. https://doi.org/10.1007/s002510050620.

[16] Valiante NM, Uhrberg M, Shilling HG, Lienert-Weidenbach K, Arnett KL, D’Andrea A, Phillips JH, Lanier LL, Parham P, N.M. Valiante, M. Uhrberg, H.G. Shilling, K. Lienert-Weidenbach, K.L. Arnett, A. D’Andrea, J.H. Phillips, L.L. Lanier, P. Parham, Functionally and structurally distinct NK cell receptor repertoires in the peripheral blood of two human donors, Immunity. 7 (1997) 739–751. http://www.ncbi.nlm.nih.gov/entrez/query.fcgi?cmd=Retrieve&db=PubMed&dopt=Citation&list_uids=9430220.

[17] N. Nemat-Gorgani, L.A. Guethlein, B.M. Henn, S.J. Norberg, J. Chiaroni, M. Sikora, L. Quintana-Murci, J.L. Mountain, P.J. Norman, P. Parham, Diversity of KIR, HLA Class I, and Their Interactions in Seven Populations of Sub-Saharan Africans, J. Immunol. 202 (2019) 2636–2647. https://doi.org/10.4049/jimmunol.1801586.

[18] C. Alicata, E. Ashouri, N. Nemat-Gorgani, L.A. Guethlein, W.M. Marin, S. Tao, L. Moretta, J.A. Hollenbach, J. Trowsdale, J.A. Traherne, A. Ghaderi, P. Parham, P.J. Norman, KIR Variation in Iranians Combines High Haplotype and Allotype Diversity With an Abundance of Functional Inhibitory Receptors, Front. Immunol. 11 (2020). https://doi.org/10.3389/fimmu.2020.00556.

[19] J. Robinson, J.A. Halliwell, J.D. Hayhurst, P. Flicek, P. Parham, S.G.E. Marsh, The IPD and IMGT/HLA database: Allele variant databases, Nucleic Acids Res. 43 (2015) D423–D431. https://doi.org/10.1093/nar/gku1161.

[20] S. Rajagopalan, Y.T. Bryceson, S.P. Kuppusamy, D.E. Geraghty, A. Van Der Meer, I. Joosten, E.O. Long, Activation of NK cells by an endocytosed receptor for soluble HLA-G, PLoS Biol. 4 (2006) 0070–0086. https://doi.org/10.1371/journal.pbio.0040009.

[21] S. Rajagopalan, E.O. Long, KIR2DL4 (CD158d): An activation receptor for HLA-G, Front. Immunol. 3 (2012) 1–6. https://doi.org/10.3389/fimmu.2012.00258.

[22] S. Kovats, E.K. Main, C. Librach, M. Stubblebine, S.J. Fisher, R. Demars, A class I antigen, HLA-G, expressed in human trophoblasts, Science (80-.). (1990). https://doi.org/10.1126/science.2326636.

[23] M.T. McMaster, C.L. Librach, Y. Zhou, K.H. Lim, M.J. Janatpour, R. DeMars, S. Kovats, C. Damsky, S.J. Fisher, Human placental HLA-G expression is restricted to differentiated cytotrophoblasts., J. Immunol. 154 (1995) 3771–8. http://www.ncbi.nlm.nih.gov/pubmed/7706718.

[24] A. Jurisicova, R.F. Casper, N.J. Maclusky, G.B. Mills, C.L. Librach, HLA-G expression during preimplantation human embryo development, Proc. Natl. Acad. Sci. U. S. A. 93 (1996) 161–165. https://doi.org/10.1073/pnas.93.1.161.

[25] M. Le Discorde, P. Moreau, P. Sabatier, J.M. Legeais, E.D. Carosella, Expression of HLA-G in Human Cornea, an Immune-Privileged Tissue, Hum. Immunol. 64 (2003) 1039–1044. https://doi.org/10.1016/j.humimm.2003.08.346.

[26] V. Cirulli, J. Zalatan, M. McMaster, R. Prinsen, D.R. Salomon, C. Ricordi, B.E. Torbett, P. Meda, L. Crisa, The class I HLA repertoire of pancreatic islets comprises the nonclassical class Ib antigen HLA-G, Diabetes. 55 (2006) 1214–1222. https://doi.org/10.2337/db05-0731.

[27] V. Mallet, A. Blaschitz, L. Crisa, C. Schmitt, S. Fournel, A. King, Y.W. Loke, G. Dohr, P. Le Bouteiller, HLA-G in the human thymus: A subpopulation of medullary epithelial but not CD83+ dendritic cells expresses HLA-G as a membrane-bound and soluble protein, Int. Immunol. 11 (1999) 889–898. https://doi.org/10.1093/intimm/11.6.889.

[28] E.A. Donadi, E.C. Castelli, A. Arnaiz-Villena, M. Roger, D. Rey, P. Moreau, Implications of the polymorphism of HLA-G on its function, regulation, evolution and disease association, Cell. Mol. Life Sci. 68 (2011) 369–395. https://doi.org/10.1007/s00018-010-0580-7.

[29] E.C. Ibrahim, S. Aractingi, Y. Allory, F. Borrini, A. Dupuy, P. Duvillard, E.D. Carosella, M.F. Avril, P. Paul, Analysis of HLA antigen expression in benign and malignant melanocytic lesions reveals that upregulation of HLA-G expression correlates with malignant transformation, high inflammatory infiltration and HLA-A1 genotype, Int. J. Cancer. 108 (2004) 243–250. https://doi.org/10.1002/ijc.11456.

[30] E.D. Carosella, B. Favier, N. Rouas-Freiss, P. Moreau, J. Lemaoult, Beyond the increasing complexity of the immunomodulatory HLA-G molecule, Blood. 111 (2008) 4862–4870. https://doi.org/10.1182/blood-2007-12-127662.

[31] B. Seliger, H. Abken, S. Ferrone, HLA-G and MIC expression in tumors and their role in anti-tumor immunity, Trends Immunol. 24 (2003) 82–87. https://doi.org/10.1016/S1471-4906(02)00039-X.

[32] R. Rizzo, D. Bortolotti, S. Bolzani, E. Fainardi, HLA-G molecules in autoimmune diseases and infections, Front. Immunol. 5 (2014) 1–11. https://doi.org/10.3389/fimmu.2014.00592.

[33] R. Rizzo, E. Fainardi, N. Rouas-Freiss, F. Morandi, The Role of HLA-Class Ib Molecules in Immune-Related Diseases, Tumors, and Infections 2016, J. Immunol. Res. 2017 (2017). https://doi.org/10.1155/2017/2309574.

[34] J.P. Goodridge, C.S. Witt, F.T. Christiansen, H.S. Warren, KIR2DL4 (CD158d) Genotype Influences Expression and Function in NK Cells, J. Immunol. 171 (2003) 1768–1774. https://doi.org/10.4049/jimmunol.171.4.1768.

[35] J.P. Goodridge, L.J. Lathbury, E. John, A.K. Charles, F.T. Christiansen, C.S. Witt, The genotype of the NK cell receptor, KIR2DL4, influences INFγ secretion by decidual natural killer cells, Mol. Hum. Reprod. 15 (2009) 489–497. https://doi.org/10.1093/molehr/gap039.

[36] I. Nowak, A. Malinowski, E. Barcz, J.R. Wilczyński, M. Wagner, E. Majorczyk, H. Motak-Pochrzęst, M. Banasik, P. Kuśnierczyk, Possible Role of HLA-G, LILRB1 and KIR2DL4 Gene Polymorphisms in Spontaneous Miscarriage, Arch. Immunol. Ther. Exp. (Warsz). 64 (2016) 505–514. https://doi.org/10.1007/s00005-016-0389-7.

[37] C.Y. Tan, Y.S. Chong, A. Loganath, Y.H. Chan, J. Ravichandran, C.G. Lee, S.S. Chong, Possible gene-gene interaction of KIR2DL4 with its cognate ligand HLA-G in modulating risk for preeclampsia, Reprod. Sci. 16 (2009) 1135–1143. https://doi.org/10.1177/1933719109342280.

[38] W. hua Yan, A. Lin, B. guo Chen, M. ying Zhou, M. zhen Dai, X. jiao Chen, L. hong Gan, M. Zhu, W. wu Shi, B. li Li, Possible roles of KIR2DL4 expression on uNK cells in human pregnancy, Am. J. Reprod. Immunol. 57 (2007) 233–242. https://doi.org/10.1111/j.1600-0897.2007.00469.x.

[39] M. Fondevila, C. Phillips, C. Santos, A. Freire Aradas, P.M. Vallone, J.M. Butler, M. V. Lareu, Á. Carracedo, Revision of the SNPforID 34-plex forensic ancestry test: Assay enhancements, standard reference sample genotypes and extended population studies, Forensic Sci. Int. Genet. 7 (2013) 63–74. https://doi.org/10.1016/j.fsigen.2012.06.007.

[40] J.K. Pritchard, M. Stephens, P. Donnelly, Inference of population structure using multilocus genotype data, Genetics. (2000).

[41] C. Posth, N. Nakatsuka, I. Lazaridis, P. Skoglund, S. Mallick, et al., Reconstructing the Deep Population History of Central and South America, Cell. 175 (2018) 1185–1197.e22. https://doi.org/10.1016/j.cell.2018.10.027.

[42] H. Li, R. Durbin, Fast and accurate short read alignment with Burrows-Wheeler transform, Bioinformatics. 25 (2009) 1754–1760. https://doi.org/10.1093/bioinformatics/btp324.

[43] E.C. Castelli, M.A. Paz, A.S. Souza, J. Ramalho, C.T. Mendes-Junior, Hla-mapper: An application to optimize the mapping of HLA sequences produced by massively parallel sequencing procedures, Hum. Immunol. 79 (2018) 678–684. https://doi.org/10.1016/j.humimm.2018.06.010.

[44] A. Souza, P. Sonon, M. Paz, L. Tokplonou, T. Lima, I. Porto, H. Andrade, N. dos Silva, L. Veiga-Castelli, M.L. Oliveira, I.A. Sadissou, J.D. Massaro, K. Moutairou, E. Donadi, A. Massougbodji, A. Garcia, M. Ibikounlé, D. Meyer, A. Sabbagh, C. Mendes-Junior, D. Courtin, E. Castelli, HLA-C Genetic Diversity and Evolutionary Insights in Two Samples From Brazil and Benin, Hla. (2020). https://doi.org/10.1111/tan.13996.

[45] T.H.A. Lima, A.S. Souza, I.O.P. Porto, M.A. Paz, L.C. Veiga-Castelli, M.L.G. Oliveira, E.A. Donadi, D. Meyer, A. Sabbagh, C.T. Mendes-Junior, E.C. Castelli, HLA-A promoter, coding, and 3’UTR sequences in a Brazilian cohort, and their evolutionary aspects, Hla. 93 (2019) 65–79. https://doi.org/10.1111/tan.13474.

[46] D.A. Benson, M. Cavanaugh, K. Clark, I. Karsch-Mizrachi, J. Ostell, K.D. Pruitt, E.W. Sayers, GenBank, Nucleic Acids Res. 46 (2018) D41–D47. https://doi.org/10.1093/nar/gkx1094.

[47] G.A. Van der Auwera, M.O. Carneiro, C. Hartl, R. Poplin, G. del Angel, A. Levy-Moonshine, T. Jordan, K. Shakir, D. Roazen, J. Thibault, E. Banks, K. V. Garimella, D. Altshuler, S. Gabriel, M.A. DePristo, From fastQ data to high-confidence variant calls: The genome analysis toolkit best practices pipeline, Curr. Protoc. Bioinforma. (2013). https://doi.org/10.1002/0471250953.bi1110s43.

[48] M.A. Depristo, E. Banks, R. Poplin, K. V. Garimella, J.R. Maguire, C. Hartl, A.A. Philippakis, G. Del Angel, M.A. Rivas, M. Hanna, A. McKenna, T.J. Fennell, A.M. Kernytsky, A.Y. Sivachenko, K. Cibulskis, S.B. Gabriel, D. Altshuler, M.J. Daly, A framework for variation discovery and genotyping using nextgeneration DNA sequencing data, Nat. Genet. 43 (2011) 491–501. https://doi.org/10.1038/ng.806.

[49] O. Delaneau, J.F. Zagury, M.R. Robinson, J.L. Marchini, E.T. Dermitzakis, Accurate, scalable and integrative haplotype estimation, Nat. Commun. 10 (2019) 24–29. https://doi.org/10.1038/s41467-019-13225-y.

[50] B.L. Browning, Y. Zhou, S.R. Browning, A One-Penny Imputed Genome from Next-Generation Reference Panels, Am. J. Hum. Genet. 103 (2018) 338–348. https://doi.org/10.1016/j.ajhg.2018.07.015.

[51] S.R. Browning, B.L. Browning, Rapid and accurate haplotype phasing and missing-data inference for whole-genome association studies by use of localized haplotype clustering, Am. J. Hum. Genet. 81 (2007) 1084–1097. https://doi.org/10.1086/521987.

[52] P. Rice, L. Longden, A. Bleasby, EMBOSS: The European Molecular Biology Open Software Suite, Trends Genet. (2000). https://doi.org/10.1016/S0168-9525(00)02024-2.

[53] L. Excoffier, G. Laval, S. Schneider, Arlequin (version 3.0): An integrated software package for population genetics data analysis, Evol. Bioinforma. 1 (2005) 117693430500100. https://doi.org/10.1177/117693430500100003.

[54] J.C. Barrett, B. Fry, J. Maller, M.J. Daly, Haploview: Analysis and visualization of LD and haplotype maps, Bioinformatics. 21 (2005) 263–265. https://doi.org/10.1093/bioinformatics/bth457.

[55] M. Faure, E.O. Long, KIR2DL4 (CD158d), an NK Cell-Activating Receptor with Inhibitory Potential, J. Immunol. 168 (2002) 6208–6214. https://doi.org/10.4049/jimmunol.168.12.6208.

[56] A. Kikuchi-Maki, S. Yusa, T.L. Catina, K.S. Campbell, KIR2DL4 Is an IL-2-Regulated NK Cell Receptor That Exhibits Limited Expression in Humans but Triggers Strong IFN-γ Production, J. Immunol. 171 (2003) 3415–3425. https://doi.org/10.4049/jimmunol.171.7.3415.

[57] J.P. Goodridge, L.J. Lathbury, N.K. Steiner, C.N. Shulse, P. Pullikotil, N.G. Seidah, C.K. Hurley, F.T. Christiansen, C.S. Witt, Three common alleles of KIR2DL4 (CD158d) encode constitutively expressed, inducible and secreted receptors in NK cells, Eur. J. Immunol. 37 (2007) 199–211. https://doi.org/10.1002/eji.200636316.

[58] W. Jiang, C. Johnson, J. Jayaraman, N. Simecek, J. Noble, M.F. Moffatt, W.O. Cookson, J. Trowsdale, J.A. Traherne, Copy number variation leads to considerable diversity for B but not A haplotypes of the human KIR genes encoding NK cell receptors, Genome Res. 22 (2012) 1845–1854. https://doi.org/10.1101/gr.137976.112.

[59] I. Nowak, E. Majorczyk, R. Ploski, D. Senitzer, J.Y. Sun, P. Kuśnierczyk, Lack of KIR2DL4 gene in a fertile Caucasian woman, Tissue Antigens. 78 (2011) 115–119. https://doi.org/10.1111/j.1399-0039.2011.01711.x.

[60] P.J. Norman, L. Abi-Rached, K. Gendzekhadze, J.A. Hammond, A.K. Moesta, D. Sharma, T. Graef, K.L. McQueen, L.A. Guethlein, C.V.F. Carrington, D. Chandanayingyong, Y.H. Chang, C. Crespí, G. Saruhan-Direskeneli, K. Hameed, G. Kamkamidze, K.A. Koram, Z. Layrisse, N. Matamoros, J. Milà, H.P. Myoung, R.M. Pitchappan, D. Dan Ramdath, M.Y. Shiau, H.A.F. Stephens, S. Struik, D. Tyan, D.H. Verity, R.W. Vaughan, R.W. Davis, P.A. Fraser, E.M. Riley, M. Ronaghi, P. Parham, Meiotic recombination generates rich diversity in NK cell receptor genes, alleles, and haplotypes, Genome Res. 19 (2009) 757–769. https://doi.org/10.1101/gr.085738.108.

[61] N. Gömez-Lozano, R. de Pablo, S. Puente, C. Vilches, Recognition of HLA-G by the NK cell receptor KIR2DL4 is not essential for human reproduction, Eur. J. Immunol. 33 (2003) 639–644. https://doi.org/10.1002/eji.200323741.

[62] S. Vendelbosch, M. de Boer, R.A.T.W. Gouw, C.K.Y. Ho, J. Geissler, W.T.N. Swelsen, M.J. Moorhouse, N.M. Lardy, D. Roos, T.K. van den Berg, T.W. Kuijpers, Extensive Variation in Gene Copy Number at the Killer Immunoglobulin-Like Receptor Locus in Humans, PLoS One. 8 (2013) 4–13. https://doi.org/10.1371/journal.pone.0067619.

[63] J. Bruijnesteijn, M.K.H. van der Wiel, N. de Groot, N. Otting, A.J.M. de Vos-Rouweler, N.M. Lardy, N.G. de Groot, R.E. Bontrop, Extensive Alternative Splicing of KIR Transcripts, Front. Immunol. 9 (2018) 2846. https://doi.org/10.3389/fimmu.2018.02846.

[64] K.L. Hershberger, R. Shyam, A. Miura, N.L. Letvin, Diversity of the Killer Cell Ig-Like Receptors of Rhesus Monkeys, J. Immunol. 166 (2001) 4380–4390. https://doi.org/10.4049/jimmunol.166.7.4380.

[65] L.A. Guethlein, L.R. Flodin, E.J. Adams, P. Parham, NK Cell Receptors of the Orangutan (Pongo pygmaeus): A Pivotal Species for Tracking the Coevolution of Killer Cell Ig-Like Receptors with MHC-C, J. Immunol. 169 (2002) 220–229. https://doi.org/10.4049/jimmunol.169.1.220.

[66] A. Auton, G.R. Abecasis, D.M. Altshuler, R.M. Durbin, et al., A global reference for human genetic variation, Nature. 526 (2015) 68–74. https://doi.org/10.1038/nature15393.

[67] J.C. Boyington, P.D. Sun, A structural perspective on MHC class I recognition by killer cell immunoglobulin-like receptors, Mol. Immunol. 38 (2002) 1007–1021. https://doi.org/10.1016/S0161-5890(02)00030-5.

[68] S. Buhler, J. Di Cristofaro, C. Frassati, A. Basire, V. Galicher, J. Chiaroni, C. Picard, High levels of molecular polymorphism at the KIR2DL4 locus in French and Congolese populations: Impact for anthropology and clinical studies, Hum. Immunol. 70 (2009) 953–959. https://doi.org/10.1016/j.humimm.2009.08.002.

[69] M.A. Gedil, N.K. Steiner, C.K. Hurley, Genomic characterization of KIR2DL4 in families and unrelated individuals reveals extensive diversity in exon and intron sequences including a common frameshift variation occurring in several alleles, Tissue Antigens. 65 (2005) 402–418. https://doi.org/10.1111/j.1399-0039.2005.00380.x.

[70] D.C. Jones, R.S. Edgar, T. Ahmad, J.R.F. Cummings, D.P. Jewell, J. Trowsdale, N.T. Young, Killer Ig-like receptor (KIR) genotype and HLA ligand combinations in ulcerative colitis susceptibility, Genes Immun. 7 (2006) 576–582. https://doi.org/10.1038/sj.gene.6364333.

[71] F.M. Zhu, K. Jiang, Q.F. Lv, J. He, L.X. Yan, Investigation of killer cell immunoglobulin-like receptor KIR2DL4 diversity by sequence-based typing in Chinese population, Tissue Antigens. 67 (2006) 214–221. https://doi.org/10.1111/j.1399-0039.2006.00562.x.

[72] K. Gendzekhadze, P.J. Norman, L. Abi-Rached, T. Graef, A.K. Moesta, Z. Layrisse, P. Parham, Coevolution of KIR2DL3 with HLA-C in a human population retaining minimal essential diversity of KIR and HLA class I ligands, Proc. Natl. Acad. Sci. U. S. A. 106 (2009) 18692–18697. https://doi.org/10.1073/pnas.0906051106.

[73] A. Goris, R. Dobosi, S. Boonen, G. Nagels, B. Dubois, KIR2DL4 (CD158d) polymorphisms and susceptibility to multiple sclerosis, J. Neuroimmunol. 210 (2009) 113–115. https://doi.org/10.1016/j.jneuroim.2009.03.001.

[74] M. Yawata, N. Yawata, M. Draghi, A.M. Little, F. Partheniou, P. Parham, Roles for HLA and KIR polymorphisms in natural killer cell repertoire selection and modulation of effector function, J. Exp. Med. 203 (2006) 633–645. https://doi.org/10.1084/jem.20051884.

[75] E.C. Castelli, J. Ramalho, I.O.P. Porto, T.H.A. Lima, L.P. Felício, A. Sabbagh, E.A. Donadi, C.T. Mendes-Junior, Insights into HLA-G genetics provided by worldwide haplotype diversity, Front. Immunol. 5 (2014). https://doi.org/10.3389/fimmu.2014.00476.

[76] P. Sonon, I. Sadissou, L. Tokplonou, K.K.G. M’po, S.S.C. Glitho, P. Agniwo, M. Ibikounlé, J.D. Massaro, A. Massougbodji, P. Moreau, A. Sabbagh, C.T. Mendes-Junior, K.A. Moutairou, E.C. Castelli, D. Courtin, E.A. Donadi, HLA-G, -E and -F regulatory and coding region variability and haplotypes in the Beninese Toffin population sample, Mol. Immunol. 104 (2018) 108–127. https://doi.org/10.1016/j.molimm.2018.08.016.

[77] D.Y.C. Brandt, V.R.C. Aguiar, B.D. Bitarello, K. Nunes, J. Goudet, D. Meyer, Mapping bias overestimates reference allele frequencies at the HLA genes in the 1000 genomes project phase I data, G3 Genes, Genomes, Genet. 5 (2015) 931–941. https://doi.org/10.1534/g3.114.015784.

[78] H. Li, P.W. Wright, M. McCullen, S.K. Anderson, Characterization of KIR intermediate promoters reveals four promoter types associated with distinct expression patterns of KIR subtypes, Genes Immun. (2016). https://doi.org/10.1038/gene.2015.56.

[79] Z. Tan, A.M. Shon, C. Ober, Evidence of balancing selection at the HLA-G promoter region, Hum. Mol. Genet. 14 (2005) 3619–3628. https://doi.org/10.1093/hmg/ddi389.

[80] E.C. Castelli, C.T. Mendes-Junior, L.C. Veiga-Castelli, M. Roger, P. Moreau, E.A. Donadi, A comprehensive study of polymorphic sites along the HLA-G gene: Implication for gene regulation and evolution, Mol. Biol. Evol. 28 (2011) 3069–3086. https://doi.org/10.1093/molbev/msr138.

[81] L. Gineau, P. Luisi, E.C. Castelli, J. Milet, D. Courtin, N. Cagnin, B. Patillon, H. Laayouni, P. Moreau, E.A. Donadi, A. Garcia, A. Sabbagh, Balancing immunity and tolerance: Genetic footprint of natural selection in the transcriptional regulatory region of HLA-G, Genes Immun. 16 (2015) 57–70. https://doi.org/10.1038/gene.2014.63.

[82] H.-I. Trompeter, N. Gómez-Lozano, S. Santourlidis, B. Eisermann, P. Wernet, C. Vilches, M. Uhrberg, Three Structurally and Functionally Divergent Kinds of Promoters Regulate Expression of Clonally Distributed Killer Cell Ig-Like Receptors (KIR), of KIR2DL4, and of KIR3DL3, J. Immunol. 174 (2005) 4135–4143. https://doi.org/10.4049/jimmunol.174.7.4135.

