## Supplementary Table 1 for "*KIR2DL4* genetic diversity in a Brazilian population sample: implications for transcription regulation and protein diversity in samples with different ancestry backgrounds"

**Table S1: The *KIR2DL4* variable sites detected in Brazilian samples from the State of São Paulo and the frequency of the reference allele.**

| Chr19 Position (hg38) | SNPid | KIR2DL4 region | IMGT/KIR relative position | Reference allele | Alternative allele(s) | Considering copy number (2n = 215) | Assuming all samples with two *KIR2DL4* copies (2n = 440) | 1000G frequency (2n=5008) | Samples presenting the reference allele (n = 220) | Notes |
|---|---|---|---|---|---|---|---|---|---|---|
| 54801760 | rs34015747 | Promoter | -1892 | T | C | 0.5814 | 0.5659 | . | 0.7818 | E |
| 54802107 | rs755822490 | Promoter | -1545 | C | A | 0.9860 | 0.9932 | . | 1.0000 | E |
| 54802354 | rs3916556 | Promoter | -1298 | C | T | 0.5814 | 0.5659 | . | 0.7818 | E |
| 54802427 | rs111803848 | Promoter | -1225 | G | T | 0.8651 | 0.8614 | . | 0.9636 | E |
| 54802632 | rs35250545 | Promoter | -1020 | C | A | 0.5814 | 0.5659 | . | 0.7818 | E |
| 54802639 | rs35160118 | Promoter | -1013 | T | C | 0.5814 | 0.5659 | . | 0.7818 | E |
| 54802679 | rs34370734 | Promoter | -973 | G | A | 0.5814 | 0.5659 | . | 0.7818 | E |
| 54802832 | rs589158 | Promoter | -820 | A | G | 0.2047 | 0.1932 | . | 0.3545 | E |
| 54803022 | rs1168390911 | Promoter | -630 | A | G | 1.0000 | 0.9977 | . | 1.0000 | E,F |
| 54803043 | rs748539788 | Promoter | -609 | T | C | 1.0000 | 0.9977 | . | 1.0000 | E,F |
| 54803145 | rs376798518 | Promoter | -507 | C | T | 0.9721 | 0.9818 | . | 1.0000 | E |
| 54803290 | rs55881846 | Promoter | -362 | G | T | 0.9953 | 0.9886 | . | 1.0000 | E |
| 54803331 | rs1175310463 | Promoter | -321 | C | CA | 0.9860 | 0.9932 | . | 1.0000 | E |
| 54803346 | rs144978425 | Promoter | -306 | C | CA | 0.2047 | 0.1932 | . | 0.3545 | E |
| 54803350 | rs36116896 | Promoter | -302 | T | G | 0.2047 | 0.1932 | . | 0.3545 | E |
| 54803379 | rs56118110 | Promoter | -273 | T | A | 0.9860 | 0.9886 | . | 1.0000 | E |
| 54803382 | rs55897924 | Promoter | -270 | C | T | 0.9953 | 0.9955 | . | 1.0000 | E |
| 54803423 | rs55808487 | Promoter | -229 | C | T | 0.9814 | 0.9705 | . | 1.0000 | A |
| 54803425 | rs56401846 | Promoter | -227 | G | A | 0.9488 | 0.9500 | . | 1.0000 | B |
| 54803437 | rs933157437 | Promoter | -215 | G | A | 1.0000 | 0.9977 | . | 1.0000 | A |
| 54803442 | rs35656676 | Promoter | -210 | G | C | 0.6791 | 0.6659 | 0.4884 | 0.8636 | B |
| 54803494 | rs17173106 | Promoter | -158 | T | C | 0.6791 | 0.6659 | 0.4882 | 0.8636 | B |
| 54803606 | rs972753648 | Promoter | -46 | G | A | 0.9953 | 0.9977 | . | 1.0000 | A |
| 54803613 | rs564804446 | Promoter | -39 | C | T | 0.9953 | 0.9977 | 0.9998 | 1.0000 | A,F |
| 54803730 | rs372376615 | Intron1 | 79 | T | C | 0.9953 | 0.9955 | . | 1.0000 | A |
| 54803852 | rs2295802 | Intron1 | 201 | C | G | 0.5442 | 0.5341 | 0.3740 | 0.7591 | B |
| 54803871 | rs604076 | Intron1 | 221 | A | C | 0.8000 | 0.7955 | . | 0.9500 | B |
| 54803917 | rs369033322 | Exon2 | 266 | G | A | 1.0000 | 0.9977 | 0.9998 | 1.0000 | A, C |
| 54803985 | rs597598 | Intron 2 | 334 | A | G | 0.2047 | 0.1932 | 0.2013 | 0.3545 | B |
| 54804045 | rs56133375 | Intron2 | 532 | G | * | 0.5953 | 0.5932 | . | 0.7864 | B |
| 54804111 | rs56136619 | Intron2 | 460 | C | T | 0.9953 | 0.9932 | . | 1.0000 | B |

| | | | | | | | | | | |
|---|---|---|---|---|---|---|---|---|---|---|
| 54804112 | rs200791941 | Intron2 | 461 | C | T | 0.8233 | 0.8386 | 0.8287 | 0.9455 | B |
| 54804161 | rs598452 | Intron2 | 544 | C | T | 0.5256 | 0.5273 | . | 0.7773 | B |
| 54804200 | rs903918210 | Intron2 | 583 | T | A | 1.0000 | 0.9932 | . | 1.0000 | A |
| 54804309 | rs113882675 | Intron2 | 692 | G | A | 0.9814 | 0.9864 | 0.9924 | 1.0000 | A |
| 54804329 | rs376787209 | Intron2 | 712 | G | C | 0.9721 | 0.9818 | 0.9830 | 1.0000 | B |
| 54804368 | rs942969021 | Intron 2 | 751 | G | C | 0.9953 | 0.9977 | . | 1.0000 | A,F |
| 54804426 | . | Intron 2 | 809 | C | T | 1.0000 | 0.9977 | . | 1.0000 | A,F |
| 54804542 | rs607149 | Intron2 | 925 | C | G | 0.2140 | 0.1977 | 0.5909 | 0.3591 | B |
| 54804576 | rs768109588 | Intron2 | 959 | A | G | 0.9907 | 0.9818 | . | 1.0000 | A |
| 54804595 | rs374093219 | Intron2 | 978 | G | A | 0.9860 | 0.9886 | 0.9870 | 1.0000 | A |
| 54804637 | rs17771905 | Intron2 | 1020 | C | T | 0.9628 | 0.9591 | . | 1.0000 | B |
| 54804643 | rs544084384 | Intron 2 | 1026 | G | A | 0.9953 | 0.9977 | 0.9996 | 1.0000 | A |
| 54804731 | rs560341968 | Intron 2 | 1114 | C | A | 1.0000 | 0.9977 | 0.9996 | 1.0000 | A,F |
| 54804874 | rs618835 | Exon 3 | 1257 | A | G | 0.7070 | 0.6909 | . | 0.9091 | B, D (Y>C) |
| 54805006 | . | Exon3 | 1389 | G | C | 1.0000 | 0.9977 | . | 1.0000 | A, D (C>W) |
| 54805109 | rs538336951 | Intron 3 | 1492 | T | A | 1.0000 | 0.9977 | 0.9988 | 1.0000 | A,F |
| 54805248 | rs2364437 | Intron3 | 1631 | G | A | 0.9721 | 0.9818 | . | 1.0000 | B |
| 54805255 | rs755023073 | Intron3 | 1638 | A | G | 0.9953 | 0.9977 | . | 1.0000 | A |
| 54805365 | rs621169 | Intron3 | 1748 | T | C | 0.7023 | 0.6886 | . | 0.9091 | B |
| 54805570 | rs2075767 | Intron3 | 1953 | C | A | 0.7953 | 0.7841 | 0.6520 | 0.9455 | B |
| 54805585 | . | Intron3 | 1968 | G | C | 0.9953 | 0.9977 | . | 1.0000 | A |
| 54805645 | rs2075768 | Intron3 | 2028 | C | T | 0.6884 | 0.6705 | 0.4926 | 0.8682 | B |
| 54805678 | rs625788 | Intron3 | 2061 | A | T | 0.8000 | 0.7955 | . | 0.9500 | B |
| 54805739 | rs531711276 | Intron3 | 2122 | G | A | 0.9953 | 0.9977 | 0.9978 | 1.0000 | A |
| 54805821 | rs148546628 | Intron3 | 2204 | G | A | 0.8698 | 0.8727 | 0.8948 | 0.9636 | B |
| 54805833 | rs1379131943 | Intron3 | 2216 | TGA | *** | 0.7023 | 0.6909 | . | 0.9091 | B |
| 54805984 | rs200485159 | Exon4 | 2367 | C | T | 0.9535 | 0.9682 | 0.9459 | 0.9955 | B, D (P>L) |
| 54806001 | rs1051454 | Exon4 | 2384 | G | A | 0.6884 | 0.6705 | 0.4840 | 0.8682 | B, D (A>T) |
| 54806015 | rs1485488219 | Exon4 | 2398 | G | C | 1.0000 | 0.9977 | . | 1.0000 | A, C |
| 54806069 | rs374413343 | Exon4 | 2452 | G | A | 0.8698 | 0.8705 | 0.8878 | 0.9636 | B, C |
| 54806095 | rs553817989 | Exon4 | 2478 | C | A | 0.9953 | 0.9977 | 0.9966 | 1.0000 | B, D (P>H) |
| 54806126 | rs577091862 | Exon4 | 2509 | G | A | 0.9953 | 0.9977 | 0.9996 | 1.0000 | A, C |
| 54806130 | rs546044168 | Exon4 | 2513 | G | A | 0.9953 | 0.9955 | 0.9996 | 1.0000 | A, D (A>N) |
| 54806157 | rs762659354 | Exon4 | 2540 | G | A | 0.9860 | 0.9932 | . | 0.9955 | A, D(G>R) |
| 54806170 | rs903729815 | Exon4 | 2553 | G | A | 0.9953 | 0.9977 | . | 1.0000 | A,F |
| 54806178 | rs562937136 | Exon4 | 2561 | G | A | 0.9953 | 0.9977 | 0.9960 | 1.0000 | B |
| 54806214 | rs1051456 | Exon4 | 2597 | C | G | 0.6884 | 0.6705 | 0.4655 | 0.8682 | B, D (Q>E) |

| | | | | | | | | | | |
|---|---|---|---|---|---|---|---|---|---|---|
| 54806367 | rs373463555 | Intron4 | 2750 | T | C | 0.8930 | 0.8886 | . | 0.9864 | B |
| 54806420 | rs923269436 | Intron4 | 2803 | C | G | 0.9953 | 0.9955 | . | 1.0000 | B |
| 54806706 | rs923501394 | Intron4 | 3089 | G | T | 0.9953 | 0.9977 | . | 1.0000 | A |
| 54806730 | rs376403025 | Intron4 | 3113 | G | C | 0.9814 | 0.9705 | . | 1.0000 | B |
| 54806736 | rs34887390 | Intron4 | 3119 | G | A | 0.9023 | 0.9000 | . | 0.9955 | A |
| 54806745 | rs657395 | Intron4 | 3128 | T | C | 0.2140 | 0.1977 | . | 0.3591 | B |
| 54806770 | rs187656554 | Intron4 | 3153 | C | T | 0.8698 | 0.8705 | . | 0.9636 | B |
| 54806816 | rs657781 | Intron4 | 3199 | T | C | 0.5907 | 0.5682 | . | 0.7682 | B |
| 54806888 | rs4806447 | Intron4 | 3271 | G | A | 0.9535 | 0.9682 | . | 0.9955 | B |
| 54806987 | rs4806564 | Intron4 | 3370 | C | T | 0.9535 | 0.9682 | . | 0.9955 | B |
| 54807013 | rs112384686 | Intron4 | 3396 | C | CTAAA | 0.6884 | 0.6705 | . | 0.8682 | A |
| 54807170 | rs78428848 | Intron4 | 3557 | C | T | 0.6884 | 0.6705 | 0.4591 | 0.8682 | B |
| 54807186 | rs374697369 | Intron4 | 3573 | C | T | 0.9721 | 0.9818 | 0.9752 | 1.0000 | B |
| 54807205 | rs659565 | Intron4 | 3592 | A | T | 0.7070 | 0.6909 | . | 0.9091 | B |
| 54807212 | rs76827991 | Intron4 | 3599 | G | C | 0.6884 | 0.6705 | 0.4591 | 0.8682 | B |
| 54807249 | rs659629 | Intron4 | 3636 | T | A | 0.1535 | 0.1455 | 0.3890 | 0.2591 | B |
| 54807462 | rs77055523 | Intron4 | 3849 | C | G | 0.9953 | 0.9955 | . | 1.0000 | A |
| 54807463 | rs113804843 | Intron4 | 3850 | G | T | 0.9395 | 0.9500 | 0.9838 | 1.0000 | A |
| 54807594 | rs112849748 | Intron4 | 3981 | C | A | 0.6884 | 0.6705 | . | 0.8682 | B |
| 54807657 | rs113418417 | Intron4 | 4044 | C | A | 0.9953 | 0.9955 | . | 1.0000 | A |
| 54807705 | rs376786209 | Intron4 | 4092 | T | C | 0.8698 | 0.8727 | . | 0.9636 | B |
| 54807723 | rs672001 | Intron4 | 4110 | A | C | 0.2140 | 0.1977 | . | 0.3591 | B |
| 54807739 | rs112947751 | Intron4 | 4126 | C | A | 0.9953 | 0.9977 | . | 1.0000 | A |
| 54807744 | rs4969516 | Intron4 | 4131 | T | C | 0.6884 | 0.6705 | . | 0.8682 | B |
| 54807818 | rs672486 | Intron4 | 4205 | T | C | 0.2140 | 0.1977 | . | 0.3591 | B |
| 54808115 | rs113064120 | Intron4 | 4502 | T | C | 1.0000 | 0.9977 | . | 1.0000 | B,F |
| 54808145 | rs78585875 | Intron4 | 4532 | G | T | 0.6884 | 0.6705 | . | 0.8682 | B |
| 54808164 | rs750734058 | Intron4 | 4551 | G | A | 1.0000 | 0.9955 | . | 1.0000 | B |
| 54808219 | rs201471326 | Intron4 | 4614 | A | AT | 0.6884 | 0.6705 | . | 0.8682 | B |
| 54808482 | rs1464775538 | Intron4 | 4877 | G | A | 0.9953 | 0.9977 | . | 1.0000 | A,F |
| 54808514 | rs796547461 | Intron4 | 4909 | G | T | 0.9953 | 0.9977 | 0.9998 | 1.0000 | A |
| 54808515 | rs373211160 | Intron4 | 4903 | G | C | 0.9628 | 0.9545 | . | 1.0000 | B |
| 54808535 | rs377627213 | Intron4 | 4923 | T | G | 0.9907 | 0.9932 | . | 1.0000 | B |
| 54808545 | rs35603706 | Intron4 | 4933 | C | T | 0.6884 | 0.6705 | . | 0.8682 | B |
| 54808591 | rs62124064 | Intron4 | 4979 | C | T | 0.9023 | 0.9000 | . | 0.9955 | A |
| 54808689 | rs770466380 | Intron4 | 5077 | A | G | 1.0000 | 0.9932 | . | 1.0000 | A |
| 54808745 | rs16985962 | Intron4 | 5133 | T | G | 0.6884 | 0.6705 | 0.5447 | 0.8682 | B |

| | | | | | | | | | | |
|---|---|---|---|---|---|---|---|---|---|---|
| 54808768 | . | Intron4 | 5156 | A | C | 1.0000 | 0.9977 | . | 1.0000 | A |
| 54808784 | rs1231830916 | Intron4 | 5172 | G | GAA, A | 0.6884 | 0.6682 | . | 0.8636 | B |
| 54808812 | rs56218903 | Intron4 | 5202 | C | A | 0.9721 | 0.9818 | . | 1.0000 | B |
| 54808824 | rs687423 | Intron4 | 5214 | G | A | 0.2140 | 0.1977 | . | 0.3591 | B |
| 54809026 | rs200439905 | Intron5 | 5416 | C | G | 0.9535 | 0.9682 | 0.9481 | 0.9955 | B |
| 54809057 | rs556397525 | Intron5 | 5447 | G | A | 0.9953 | 0.9977 | 0.9996 | 1.0000 | A |
| 54809212 | rs600898 | Intron5 | 5602 | A | C | 0.7070 | 0.6909 | . | 0.9091 | B |
| 54809237 | rs76869409 | Intron5 | 5627 | A | G | 0.6884 | 0.6705 | 0.4433 | 0.8682 | B |
| 54809324 | rs10500318 | Intron5 | 5714 | G | A | 0.9023 | 0.9000 | 0.9060 | 0.9955 | A |
| 54809414 | rs35579530 | Intron5 | 5804 | C | T | 0.6884 | 0.6705 | 0.4878 | 0.8682 | B |
| 54809472 | rs592645 | Intron5 | 5862 | T | A | 0.8000 | 0.7955 | . | 0.9500 | B |
| 54809529 | rs564396483 | Intron5 | 5919 | C | G | 0.9953 | 0.9977 | 0.9946 | 1.0000 | A |
| 54809598 | rs1015105183 | Intron5 | 5988 | T | C | 1.0000 | 0.9955 | . | 1.0000 | A |
| 54809639 | rs992444885 | Intron5 | 6029 | C | T | 1.0000 | 0.9977 | . | 1.0000 | A,F |
| 54809643 | rs112006653 | Intron5 | 6033 | G | A | 0.9953 | 0.9955 | . | 1.0000 | A |
| 54809822 | rs111735328 | Intron5 | 6212 | A | G | 1.0000 | 0.9932 | 0.9858 | 1.0000 | A |
| 54809941 | rs604077 | Intron5 | 6331 | T | C | 0.3488 | 0.3295 | 0.3385 | 0.5364 | B |
| 54809968 | rs568087247 | Intron5 | 6358 | C | G | 0.9860 | 0.9932 | 0.9998 | 0.9955 | A |
| 54810058 | rs16986024 | Intron5 | 6448 | G | A | 0.6884 | 0.6705 | 0.4948 | 0.8682 | B |
| 54810105 | rs1473785473 | Intron5 | 6495 | A | G | 0.9953 | 0.9977 | . | 1.0000 | A,F |
| 54810126 | rs138956580 | Intron5 | 6517 | GT | G | 0.8698 | 0.8727 | . | 0.9636 | B |
| 54810142 | rs604999 | Intron5 | 6532 | A | G | 0.1535 | 0.1455 | . | 0.2591 | B |
| 54810151 | rs185767446 | Intron5 | 6541 | C | T | 0.8698 | 0.8727 | 0.9199 | 0.9636 | B |
| 54810285 | rs199888594 | Intron5 | 6675 | A | AT, ATT | 0.8698 | 0.8682 | 0.9097 | 0.9636 | A |
| 54810298 | rs145370691 | Intron5 | 6688 | A | AT | 0.7488 | 0.7636 | . | 0.9409 | B |
| 54810324 | rs1417253702 | Intron5 | 6714 | T | C | 1.0000 | 0.9977 | . | 1.0000 | A |
| 54810401 | rs616496 | Intron5 | 6791 | A | C | 0.8000 | 0.7955 | . | 0.9500 | B |
| 54810518 | rs544859823 | Intron5 | 6908 | G | GA | 1.0000 | 0.9932 | 0.9848 | 1.0000 | A |
| 54810630 | rs371261951 | Intron5 | 7020 | G | C | 0.8651 | 0.8705 | 0.9271 | 0.9636 | B |
| 54810646 | rs608316 | Intron5 | 7036 | C | G | 0.6698 | 0.6614 | 0.9103 | 0.8864 | B |
| 54810647 | rs529464965 | Intron5 | 7037 | A | G | 0.8698 | 0.8727 | 0.9271 | 0.9636 | B |
| 54810672 | rs3865505 | Intron5 | 7062 | G | A | 0.9535 | 0.9682 | . | 0.9955 | B |
| 54810680 | rs113577571 | Intron5 | 7070 | A | G | 0.9953 | 0.9977 | 0.9766 | 1.0000 | A |
| 54810689 | rs608368 | Intron5 | 7079 | A | T | 0.2186 | 0.2045 | . | 0.3682 | B |
| 54810714 | rs617872 | Intron5 | 7104 | C | G | 0.2605 | 0.2341 | . | 0.4136 | B |
| 54810724 | rs3865506 | Intron5 | 7114 | C | T | 0.9535 | 0.9659 | . | 0.9955 | B |
| 54810735 | rs617907 | Intron5 | 7125 | T | C | 0.6233 | 0.6341 | . | 0.8591 | B |

| | | | | | | | | | | |
|---|---|---|---|---|---|---|---|---|---|---|
| 54810746 | rs17173113 | Intron5 | 7136 | A | C | 0.8651 | 0.8705 | . | 0.9636 | B |
| 54810921 | rs3865507 | Intron5 | 7311 | C | A | 0.9535 | 0.9659 | 0.9409 | 0.9955 | B |
| 54810968 | rs618871 | Intron5 | 7358 | G | C | 0.8000 | 0.7955 | 0.9998 | 0.9500 | B |
| 54811235 | rs3786855 | Intron5 | 7625 | T | A | 0.7953 | 0.7818 | 0.6312 | 0.9409 | B |
| 54811247 | rs111490149 | Intron5 | 7637 | A | T | 1.0000 | 0.9932 | 0.9836 | 1.0000 | A |
| 54811269 | rs630497 | Intron5 | 7659 | G | A | 0.6186 | 0.6318 | . | 0.8591 | B |
| 54811291 | rs3786857 | Intron5 | 7681 | G | A | 0.7442 | 0.7341 | 0.5451 | 0.9182 | B |
| 54811324 | rs112954606 | Intron5 | 7714 | G | C | 1.0000 | 0.9932 | 0.9836 | 1.0000 | A |
| 54811342 | rs377528700 | Intron5 | 7732 | G | C | 0.9953 | 0.9977 | 0.9930 | 1.0000 | A |
| 54811353 | rs1294892223 | Intron5 | 7743 | G | A | 1.0000 | 0.9977 | . | 1.0000 | A |
| 54811364 | rs368909099 | Intron5 | 7754 | C | T | 0.9814 | 0.9795 | 0.9545 | 1.0000 | B |
| 54811419 | rs201457831 | Intron5 | 7809 | C | A | 0.9535 | 0.9659 | 0.9447 | 0.9955 | B |
| 54811442 | rs1344820062 | Intron5 | 7832 | G | A | 1.0000 | 0.9977 | . | 1.0000 | A |
| 54811520 | rs631717 | Intron5 | 7910 | G | A | 0.4884 | 0.4682 | . | 0.7000 | A |
| 54811534 | rs1380628341 | Intron5 | 7924 | C | T | 0.9953 | 0.9977 | . | 1.0000 | A |
| 54811574 | rs112264748 | Intron5 | 7964 | T | A | 1.0000 | 0.9932 | 0.9858 | 1.0000 | A |
| 54811602 | rs632126 | Intron5 | 7992 | A | G | 0.8000 | 0.7886 | . | 0.9500 | B |
| 54811619 | rs56136222 | Intron5 | 8009 | G | C | 1.0000 | 0.9932 | . | 1.0000 | B |
| 54811638 | rs17771961 | Intron5 | 8028 | C | G | 0.8698 | 0.8727 | 0.9050 | 0.9636 | B |
| 54811872 | rs938247389 | Intron5 | 8262 | G | A | 1.0000 | 0.9977 | . | 1.0000 | A,F |
| 54811879 | rs1302581073 | Intron5 | 8269 | A | T | 1.0000 | 0.9977 | . | 1.0000 | A,F |
| 54812145 | rs77611365 | Intron5 | 8535 | G | C | 0.6372 | 0.6364 | . | 0.8455 | B |
| 54812181 | rs371450241 | Intron5 | 8571 | G | T | 0.9488 | 0.9523 | . | 1.0000 | B |
| 54812575 | rs3865508 | Intron5 | 8965 | T | G | 0.6419 | 0.6386 | . | 0.8500 | B |
| 54812586 | rs3865509 | Intron5 | 8976 | C | G | 0.9535 | 0.9659 | . | 0.9955 | B |
| 54812599 | rs1219302608 | Intron5 | 8989 | A | G | 1.0000 | 0.9977 | . | 1.0000 | A,F |
| 54812753 | rs1412718154 | Intron5 | 9143 | C | T | 1.0000 | 0.9977 | . | 1.0000 | A |
| 54812771 | rs1197427000 | Intron5 | 9161 | G | C | 1.0000 | 0.9977 | . | 1.0000 | A,F |
| 54812784 | rs3865510 | Intron5 | 9174 | C | A | 0.6419 | 0.6318 | . | 0.8409 | B |
| 54812863 | rs11375633 | Intron5 | 9254 | A | AC | 0.2140 | 0.2000 | . | 0.3591 | B |
| 54812896 | rs537009792 | Intron5 | 9287 | G | A | 1.0000 | 0.9977 | 0.9996 | 1.0000 | A,F |
| 54812975 | rs55736818 | Intron5 | 9366 | G | A | 1.0000 | 0.9977 | . | 1.0000 | A |
| 54813019 | rs648689 | Intron5 | 9410 | T | C | 0.4419 | 0.4295 | 0.5821 | 0.6636 | B |
| 54813111 | rs649117 | Intron5 | 9502 | C | T | 0.4419 | 0.4295 | 0.5853 | 0.6636 | B |
| 54813158 | rs192263392 | Exon6 | 9549 | A | G | 0.9953 | 0.9977 | 0.9996 | 1.0000 | B,F |
| 54813180 | rs649216 | Exon6 | 9571 | T | C | 0.4419 | 0.4295 | 0.4093 | 0.6636 | B, C |
| 54813219 | rs11371265 | Exon6 | 9620 | C | CA | 0.4512 | 0.4386 | . | 0.6818 | B, D (truncated) |

| | | | | | | | | | | |
|---|---|---|---|---|---|---|---|---|---|---|
| 54813335 | rs755387785 | Intron6 | 9726 | G | T | 1.0000 | 0.9977 | . | 1.0000 | A |
| 54813377 | rs660437 | Intron6 | 9769 | C | A | 0.8000 | 0.7977 | . | 0.9500 | B |
| 54813405 | rs660773 | Intron6 | 9797 | G | A | 0.4419 | 0.4295 | 0.5186 | 0.6636 | B |
| 54813482 | . | Intron6 | 9874 | A | G | 1.0000 | 0.9977 | . | 1.0000 | A |
| 54813668 | rs652671 | Intron6 | 10060 | T | C | 0.8000 | 0.7909 | 0.9996 | 0.9500 | B |
| 54813679 | rs372293579 | Intron6 | 10071 | G | A | 1.0000 | 0.9977 | . | 1.0000 | A,F |
| 54813692 | rs201370666 | Exon7 | 10084 | G | A | 0.9953 | 0.9977 | 0.9984 | 1.0000 | A, D(A>T) |
| 54813785 | rs653140 | Intron7 | 10177 | G | C | 0.4419 | 0.4295 | . | 0.6636 | B |
| 54813786 | rs653142 | Intron7 | 10178 | A | T | 0.4419 | 0.4295 | . | 0.6636 | B |
| 54813812 | rs751213391 | Intron7 | 10204 | C | T | 0.9953 | 0.9977 | . | 1.0000 | A |
| 54813817 | rs1445952284 | Intron7 | 10209 | T | C | 0.9953 | 0.9977 | . | 1.0000 | A,F |
| 54813863 | rs56184317 | Exon8 | 10255 | C | G | 1.0000 | 0.9932 | . | 1.0000 | A, D (Q>E) |
| 54813995 | rs35946789 | Exon8 | 10387 | G | C | 0.9860 | 0.9886 | . | 1.0000 | A, D (A>P) |
| 54813996 | rs374021043 | Exon8 | 10388 | C | T | 0.9953 | 0.9955 | 0.9890 | 1.0000 | A, D (A>V, L) |
| 54814000 | rs1051457 | Exon8 | 10392 | G | A | 0.8093 | 0.8091 | 0.7722 | 0.9545 | B, C |
| 54814076 | rs200678651 | Exon8 | 10468 | C | T | 1.0000 | 0.9977 | 0.9966 | 1.0000 | B |
| 54814116 | rs370869656 | 3'UTR | 10508 | G | A | 0.9860 | 0.9909 | 0.9996 | 1.0000 | A |
| 54814169 | rs376917646 | 3'UTR | 10561 | G | C | 0.9628 | 0.9545 | . | 1.0000 | B |
| 54814205 | . | 3'UTR | 10597 | T | C | 0.9953 | 0.9977 | . | 1.0000 | A |
| 54814353 | rs1159738890 | 3'UTR | 10745 | GGCT | G | 0.9953 | 0.9977 | . | 1.0000 | A,F |
| 54814361 | rs34785252 | 3'UTR | 10753 | C | A | 0.8093 | 0.8114 | 0.7204 | 0.9545 | B |
| 54814366 | rs374254587 | 3'UTR | 10758 | C | G | 0.9721 | 0.9818 | . | 1.0000 | B |
| 54814422 | rs1443295872 | 3'UTR | 10814 | A | G | 1.0000 | 0.9977 | . | 1.0000 | A,F |

* The alternative allele is GGGGAGTCTCTCATGAACTAGTAAGAGGAGATCCT.

*** The alternative allele are TGAGAGA, TGAGAGAGA, TGAGAGAGAGA

Notes: (A) This variant is not described in the IPD-IMGT/KIR database; (B) This variant is described in the IPD-IMGT/KIR database; (C) synonymous exchange in exon; (D) non-synonymous exchange in exon; (E) region not tracked by the IPD-IMGT/KIR database; (F) unphased singleton . A dot (.) in the 1000Genomes (1000g) data indicates that this variant is not included in the 1000g dataset.
