## Supplementary Table 2 for "*KIR2DL4* genetic diversity in a Brazilian population sample: implications for transcription regulation and protein diversity in samples with different ancestry backgrounds"

Table S2: Comparison of the hla-mapper workflow and PING in 25 samples from Brazil

| Sample | hla-mapper [a] (CDS) | PING [b] | Notes |
|---|---|---|---|
| 01 | KIR2DL4*00501, KIR2DL4*00801 | KIR2DL4*00501, KIR2DL4*00801 | identical results |
| 02 | KIR2DL4*00501, KIR2DL4*00501 | KIR2DL4*00501, new.allele.near (*00502) | PING failed rs11371265, all other results are identical |
| 03 | KIR2DL4*00102, KIR2DL4*00801 | KIR2DL4*00102, KIR2DL4*00801 | identical results |
| 04 | KIR2DL4*00501, KIR2DL4*00801 | KIR2DL4*00501, KIR2DL4*00801 | identical results |
| 05 | KIR2DL4*00102, KIR2DL4*00501 | KIR2DL4*00102, new.allele.near (*00502) | PING failed rs11371265, all other results are identical |
| 06 | KIR2DL4*00102, KIR2DL4*00801 | KIR2DL4*00102, KIR2DL4*00801 | identical results |
| 07 | KIR2DL4*00801, KIR2DL4*01101 | KIR2DL4*00801, KIR2DL4*01101 | identical results |
| 08 | KIR2DL4*00103, KIR2DL4*00602 | KIR2DL4*00103, KIR2DL4*00602 | identical results |
| 09 | KIR2DL4*00102, KIR2DL4*00501 | KIR2DL4*00102, new.allele.near (*00502) | PING failed rs11371265, all other results are identical |
| 10 | KIR2DL4*00801, KIR2DL4*00801 | KIR2DL4*00801, KIR2DL4*00801 | identical results |
| 11 | KIR2DL4*00501, KIR2DL4*00801 | KIR2DL4*00501, KIR2DL4*00801 | identical results |
| 12 | KIR2DL4*00501, KIR2DL4*00802 | KIR2DL4*00501, KIR2DL4*00802 | identical results |
| 13 | KIR2DL4*00501, KIR2DL4*00501 | KIR2DL4*00501, KIR2DL4*00501 | identical results |
| 14 | KIR2DL4*00501, KIR2DL4*01101 | KIR2DL4*00501, KIR2DL4*01101 | identical results |
| 15 | KIR2DL4*00802, KIR2DL4*01101 | KIR2DL4*00802, KIR2DL4*01101 | identical results |
| 16 | KIR2DL4*00102, KIR2DL4*01101 | KIR2DL4*00102, KIR2DL4*01101 | identical results |
| 17 | KIR2DL4*00501, KIR2DL4*00801 | KIR2DL4*00501, KIR2DL4*00801 | identical results |
| 18 | KIR2DL4*00802, KIR2DL4*01101 | KIR2DL4*00802, KIR2DL4*01101 | identical results |
| 19 | KIR2DL4*00501, KIR2DL4*00801 | KIR2DL4*00501, KIR2DL4*00801 | identical results |
| 20 | KIR2DL4*00102, KIR2DL4*00102 | KIR2DL4*00102, new.allele.near (*00103) | PING failed rs11371265, all other results are identical |
| 21 | KIR2DL4*00103, KIR2DL4*00501 | KIR2DL4*00103, new.allele.near (*00502) | PING failed rs11371265, all other results are identical |
| 22 | KIR2DL4*00102, KIR2DL4*00801 | KIR2DL4*00102, KIR2DL4*00801 | identical results |
| 23 | KIR2DL4*00102, KIR2DL4*00501 | KIR2DL4*00102, new.allele.near (*00502) | PING failed rs11371265, all other results are identical |
| 24 | KIR2DL4*00501, KIR2DL4*00801 | KIR2DL4*00501, KIR2DL4*00801 | identical results |
| 25 | KIR2DL4*01101, new.snp | KIR2DL4*01101, new.snp | identical results |

a) hla-mapper workflow: hla-mapper for read alignment, vcfx for variant refinement, GATK HaplotypeCaller for genotype calls, phasex for haplotype estimation. b) The first version of PING, downloaded from https://hollenbachlab.ucsf.edu/ping
